# A chemical molecule promotes Trop2^+^ biliary duct organoids differentiation into insulin-secreting cells

**DOI:** 10.1101/2022.03.10.483710

**Authors:** Miao Liu, Wen Yu, Yannan Wang, Bingru Yan, Jie Shang, Congyi Zhang, Huan Wu, Sheng Tai, Liang Jin, Chun-Bo Teng

**Affiliations:** Molecular Cell Biology Laboratory, Heilongjiang Key Laboratory of Plant Bioactive Substance Biosynthesis and Utilization, College of Life Science, Northeast Forestry University, Harbin, 150040, China; Department of Hepatobiliary and Pancreatic Surgery, The Second Affiliated Hospital of Harbin Medical University, Harbin, 150086, China; State Key Laboratory of Natural Medicines, Jiangsu Key Laboratory of Druggability of Biopharmaceuticals, School of Life Science and Technology, China Pharmaceutical University, Nanjing, 210009, China

**Author notes:** **Corresponding authors:** Dr. Chun-Bo Teng, College of Life Sciences, Northeast Forestry University, No. 26, Hexing Road, Xiangfang District, Harbin, Heilongjiang province, China, Dr. Liang Jin, State Key Laboratory of Natural Medicines, Jiangsu Key Laboratory of Druggability of Biopharmaceuticals, School of Life Science and Technology, China Pharmaceutical University, 24 Tongjiaxiang, Nanjing, Jiangsu province, China, Dr. Sheng Tai, Department of Hepatobiliary and Pancreatic Surgery, The Second Affiliated Hospital of Harbin Medical University, No. 246, Xuefu Road, Nangang District, Harbin, Heilongjiang province, China. Miao Liu and Wen Yu contributed equally to this work.

**Keywords:** Trop2, extrahepatic bile duct, pancreatic duct, organoids, BMP7, chemical small molecule, insulin-secreting cell, diabetes

## Abstract

Cell replacement therapy is a hopeful strategy for diabetes treatment, whereas lack of pancreas donors leads this strategy to a dilemma. Here, we demonstrate that Trop2 is a useful marker to enrich the progenitors with a long-term organoid formation capability from extrahepatic bile ducts in both mouse and human. Based on BMP7 initiation, the Trop2-positive mouse extrahepatic bile duct organoids (mBDOs) could be induced to produce 8.49 ± 0.45% of insulin-secreting cells. We further screened out a chemical small molecule TLY142, which significantly increased the proportion of insulin-secreting cells to 19.9 ± 0.62% and 25.50 ± 4.82% in induced mouse and human extrahepatic bile duct organoids (hBDOs) *in vitro*, and transplantation of these differentiated organoids into STZ-induced diabetic nude mice remarkably improved their blood glucose and enhanced glucose sensitivity. This study provides a novel strategy that using small molecules promotes biliary duct organoids differentiation into β cells to cure diabetes.

**Graphic abstract:** 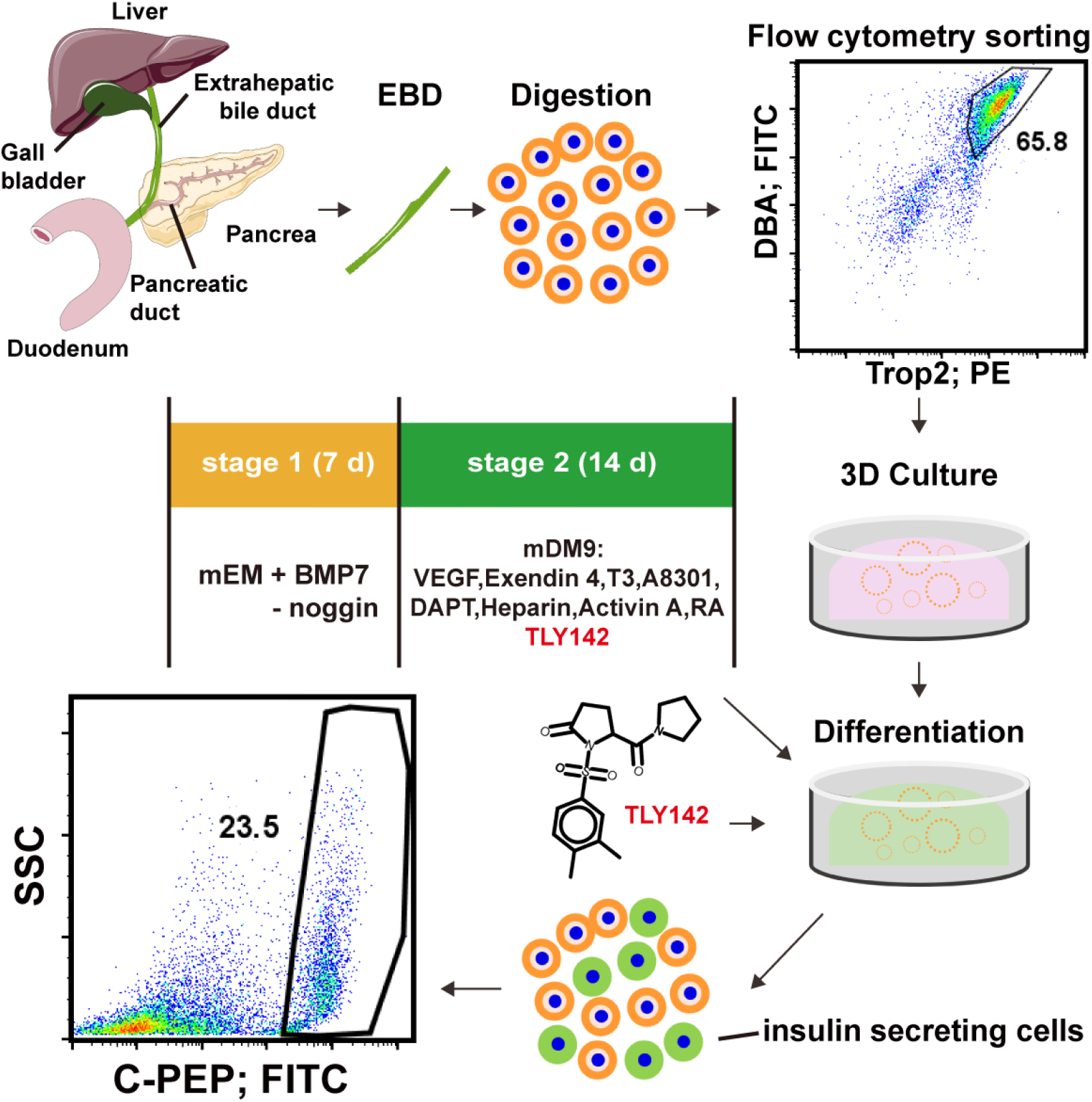

## Introduction

Cell replacement therapy is a hopeful strategy for diabetes patients with seriously destroyed beta cells to recover their healthy life (Nijhoff et al., 2016; Rickels and Robertson, 2019; Shapiro et al., 2000; Shapiro et al., 2017). Multipotent adult stem/progenitors are the potential cell source of *in vitro* differentiated beta cells (Smukler et al., 2011; Wang et al., 2020; Wang et al., 2016). Several lines of evidence demonstrate that there exist progenitor cells capable of differentiating into insulin-secreting cells in adult pancreatic ducts (Huch et al., 2013; Inada et al., 2008; Loomans et al., 2018). However, for patients with a need for cell transplantation, lack of donor organs and immune-rejection of the xenogenous cells still require to overcome.

Recent studies demonstrated that the progenitors in the biliary epithelium taken from patients could be expanded into organoids to repair the damaged biliary duct system (Sampaziotis et al., 2017; Sampaziotis and Muraro, 2021). These findings highlight a pathway to repair the damaged gastrointestinal organs using progenitors taken from patients themselves. Intriguingly, insulin-producing cells could be found in extrahepatic biliary duct epithelia and in peribiliary gland, indicating there exists progenitor cells with pancreatic differentiation potential in biliary ducts (Cardinale et al., 2011; Cardinale et al., 2012; Dutton et al., 2007). Studies on the embryonic development indicate that both the pancreatic ducts and biliary ducts initiate from the same population of progenitors situated in the boundary of mid-hind primordial gut tube at E10.5 day in mouse (Cardinale et al., 2012; Spence et al., 2009). Moreover, Ngn3-positive progenitor cells also exist in biliary ducts, and these progenitor cells present similar endocrine differentiated potential (Cardinale et al., 2011; Gribben et al., 2021). Therefore, we proposed that, enrichment and characterization of this population of progenitors in biliary ducts will be helpful for the usage of autologous biliary duct progenitor-derived organoids as islet surrogate.

Single-cell transcriptome analysis reveals that the epithelial cells are heterogenous both in pancreatic ducts or biliary ducts (Aizarani et al., 2019; Hendley et al., 2021; Qadir et al., 2020); however, which population of cells in biliary ducts can be propagated for a long term and differentiated into insulin-secreting cells is elusive. Tumor-associated calcium signal transducer 2 (Trop2) is a type I transmembrane glycoprotein, which is initially found to be highly expressed in trophoblasts and in epithelial cancers (Fornaro et al., 1995; Lipinski et al., 1981; Trerotola et al., 2013). It has been found that Trop2 represents a subpopulation of prostate progenitor cells with sphere-forming and organoid-forming capacity, which could regenerate tubular structures *in vivo* (Goldstein et al., 2008; Guo et al., 2020). Otherwise, Trop2 is highly expressed in liver oval cells with regeneration capability, and could grow into bipotent liver organoids (Aizarani et al., 2019; Okabe et al., 2009). Nevertheless, whether Trop2 marks a population of progenitors with insulin-secreting cell potential in biliary ducts is still not known.

Another puzzle in the usage of ductal progenitors is a rather low efficiency of beta-cell differentiation in both *in vivo* and *in vitro*. Loomans CJM et al. recently revealed that less than 2 percent of insulin-positive cells were obtained from the induced human pancreatic Epcam^+^ organoids (Loomans et al., 2018). In the past decades, multiple cytokines and signaling inhibitors have been explored to increase beta-cell differentiation, including fibroblast growth factor 10 (FGF10), nicotinamide, retinoic acid (RA), and exendin 4 have been demonstrated to be capable of increasing insulin expression in cultured embryonic stem cells (ES) or adult stem cells (Kassem et al., 2016; Mfopou et al., 2010; Oström et al., 2008; Pagliuca et al., 2014; Vaca et al., 2008), whereas these molecules usually promote beta-cell differentiation after the determination of progenitor to endocrine fate. Bone morphogenetic protein 7 (BMP7) is a member of transform growth factor β (TGFβ), which is proven to promote liver regeneration in partial hepatectomy mouse model (Sugimoto et al., 2007). Recently, Klein revealed that the recombinant human BMP7 plays a role in ductal to endocrine progenitor transition (Klein et al., 2015; Qadir et al., 2018), indicating it may function in the endocrine differentiation of ductal progenitor-derived organoids. Further, a higher proportion of beta cells differentiated from the endocrine-transited ductal progenitors is still required for diabetes therapy.

Chemical approach has been demonstrated to be a useful tool in facilitating the differentiation of stem/progenitor cells (Li et al., 2013; Zhu et al., 2011). Specifically in pluripotent stem cell field, chemical small molecules have been becoming powerful tools to increase a high efficiency of hepatocyte differentiation from ES or induced pluripotent stem cells (iPS) (Shan et al., 2013; Tasnim et al., 2015). Except those known activators and inhibitors of cell signaling pathways in pancreas development, there is still lack of definite chemical molecules to interrogate the unknown pathway in enhancement of beta-cell differentiation. In this study, we identified Trop2 as a novel marker enriching multipotent progenitors from the extrahepatic bile duct as well as the pancreatic main duct. We further screened out a chemical molecule which could promote mouse and human biliary duct progenitor-derived organoids differentiation into glucose-responsive beta-like cells, and tested their potential usage in diabetes therapy.

## Results

### The organoids from extrahepatic bile ducts share a similar gene expression signature with those from pancreatic ducts in mouse

To seek the available source of progenitors with long-term self-renewal and beta-cell differentiation potential, we first established mouse extrahepatic bile duct organoids (mBDOs) and pancreatic duct organoids (mPDOs), respectively, which have been considered grown from the same population of progenitors reside in the embryonic endoderm epithelium (Cardinale et al., 2011; Cardinale et al., 2012). Briefly, the epithelial cells were obtained via enzyme digestion of the mechanically stripped mouse extrahepatic bile ducts (mEBD) and pancreatic ducts (mPD), and then were embedded into Matrigel and cultured in the mouse expansion media (mEM) for 3D organoid formation (Figure 1A). On the third day, typical lumen-like structures could be observed in both mBDOs and mPDOs culture, and they gradually expanded to increase cell density with the extension of culture time (7-14 d). As anticipated, mBDOs and mPDOs have similar morphological changes (Figure 1B). The fully growing organoids in Matrigel could be passaged over 20 generation (Figure S1A), proving that the mBDOs and mPDOs had similar long-term proliferative capability. Immunostaining results showed that both mBDOs and mPDOs expressed biliary/pancreatic progenitor marker including sex determining region Y-box 9 (SOX9) and pancreatic and duodenal homeobox 1 (PDX1), development-related genes such as hepatocyte nuclear factor 1 beta (HNF1B) and hepatocyte nuclear factor 4 alpha (HNF4A), and epithelium-specific genes including epithelial cell adhesion molecule (EpCAM) and calcium-dependent cell adhesion molecule (E-cadherin) (Figure 1C, 1D, S1B and S1C).

**Figure 1.**
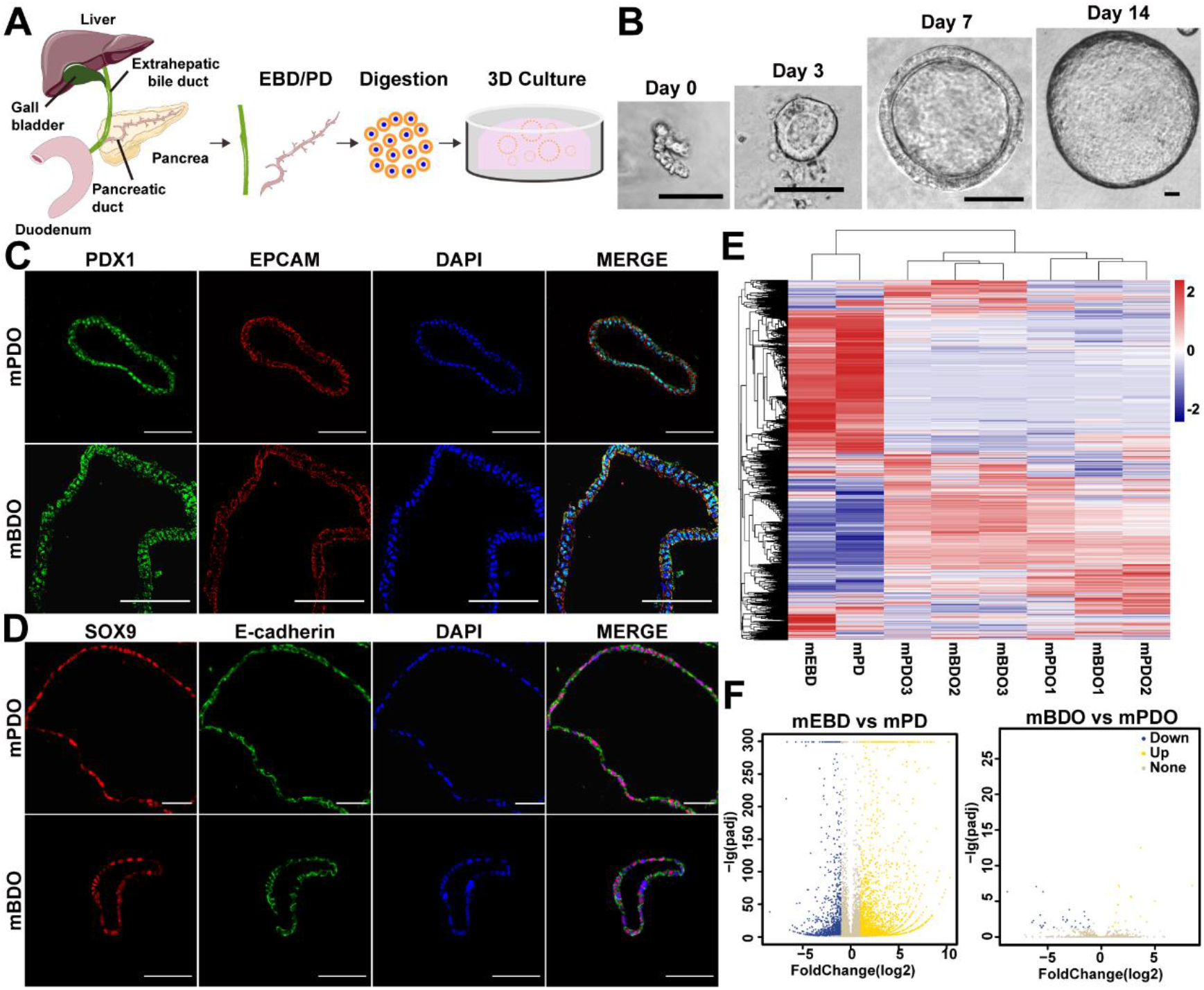
The organoids grown from mouse extrahepatic bile duct (mEBD) and pancreatic duct (mPD) have similar gene expression signature. **(A)** Schematic representation of the organoid preparation from mouse extrahepatic bile duct (mBDOs) and pancreatic duct (mPDOs). **(B)** Serial images of isolated cells and growing organoids. Scale bar, 50 μm. **(C)** and **(D)** mPDOs and mBDOs were stained with the primary antibodies against EPCAM, E-cadherin, SOX9, PDX1. Scale bar, 50 μm. **(E)** Expression heatmap of differentially expressed genes between mEBD, mPD, mBDOs and mPDOs. Blue, downregulated. Red, upregulated. **(F)** Volcano plot represents the comparation between mEBD vs mPD and mBDO vs mPDO gene signatures. X axis shows fold change (in log2), and Y axis shows adjusted p value (in -log10). Each dots represents a gene. Up-regulated genes are in yellow and down-regulated genes are in blue.

To further clarify their characteristics, we profiled mEBD, mPD, and primarily cultured mBDOs and mPDOs via a high-throughput RNA sequencing (RNA-seq). Gene expression profiling analysis showed that mBDOs and mPDOs had a more similar gene expression signature compared with their primarily isolated mEBD and mPD (Figure 1E). Analysis of the differentially expressed genes between organoids and their corresponding primary epithelial cells further confirmed the similarity between the two kinds of organoids (Figure 1F). These results indicate that mBDOs share a similar gene expression signature with mPDOs.

### Trop2 can be used as a marker to enrich progenitors capable of forming long-term cultured mBDOs from mEBD as well as mPDOs from mPD

To further confirm the similarity between mBDOs and mPDOs, we tried to find a common cell-surface marker to enrich their progenitor cells from mEBD and mPD. Based on the RNA-seq data, we looked for the up-regulated genes in both mBDOs and mPDOs compared with their primary epithelial cells, and found 16 overlapping genes (Figure 2A). Among the overlapping genes between mBDOs vs mEBD and mPDOs vs mPD, *Trop2* was found at the forefront (Figure 2B). Immunostaining of mouse extrahepatic bile duct and pancreas sections showed that Trop2 was highly expressed in mEBD and mPD (Figure 2C and S2A), but low in pancreatic acinar cells and even no expression in islets (Figure S2B). Meanwhile, flow cytometry (FCM) showed that most of mEBD and mPD epithelial cells (DBA^+^) were Trop2 positive (Trop2^+^) (Figure 2D). In the sorted cells using Trop2^+^DBA^+^ by flow cytometry, 10.50 ± 2.86% of cells from mEBD and 9.60 ± 4.94% of cells from mPD were capable of forming organoids (Figure 2E), and these mBDOs or mPDOs could be long-term passaged (Figure S2C). Next, we characterized the organoids grown from Trop2^+^ cells via immunostaining and flow cytometry. The results demonstrated that Trop2^+^ cell-derived organoids were still Trop2-positive (Figure S2D and S2E), suggesting that Trop2 can be used as a marker to enrich the progenitors with the long-term organoid formation capability from mEBD or mPD.

**Figure 2.**
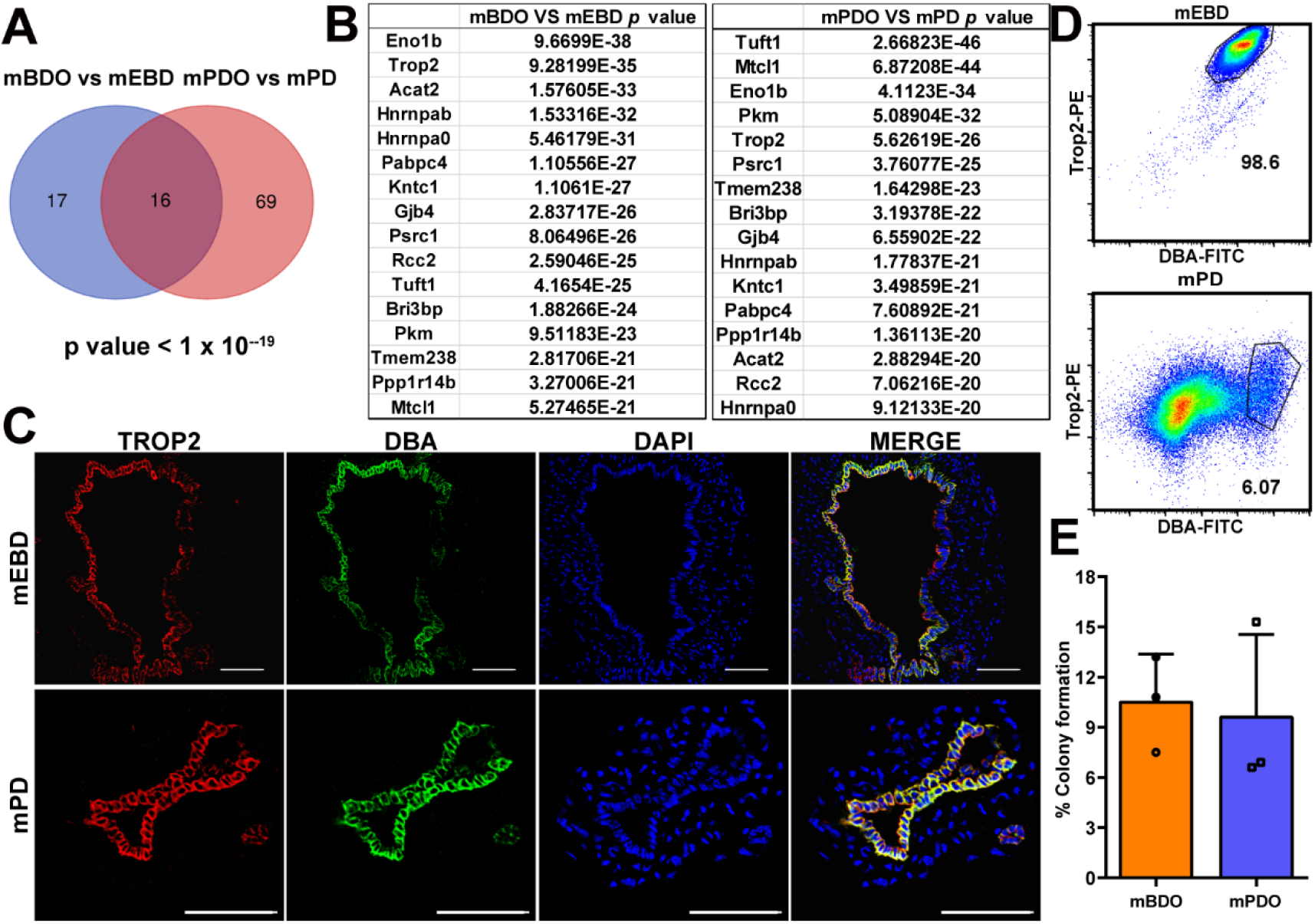
The Trop2^+^ epithelial cells in mEBD and mPD contain progenitors with organoid forming capacity. **(A)** Venn diagrams show the overlap of genes significantly up-regulated in mBDOs or mPDOs relative to mEBD or mPD. **(B)** The overlap of genes in mBDOs vs mEBD and mPDOs vs mPD. **(C)** Immunofluorescence staining of Trop2 in mEBD and mPD epithelium (DBA^+^). Scale bar, 50 μm. **(D)** Gating scheme for the isolation of Trop2^+^ cells from mEBD or mPD epithelia. **(E)** The organoid formation efficiency in Trop2^+^ DBA^+^ fraction of cells. n=3, 1000 cells were seeded each well.

### Trop2^+^ biliary organoids have a similar pancreatic differentiation potential as the pancreatic organoids in mouse

We had clarified the similar gene expression signature between mBDOs and mPDOs, and then tried to explore whether mBDOs have a similar differentiation potential as mPDOs. The previous studies on pancreatic duct organoids showed a rather low beta-cell differentiation (Huch et al., 2013; Loomans et al., 2018). It has been reported that TGF-β family member BMP7 plays a role in turning ductal epithelia to endocrine fate (Klein et al., 2015; Qadir et al., 2018), and several cytokines and inhibitors promote the differentiation of pancreatic progenitor cells (Kassem et al., 2016; Mfopou et al., 2010; Oström et al., 2008; Pagliuca et al., 2014; Vaca et al., 2008). Accordingly, we designed a 2-stage differentiation scheme for Trop2^+^ progenitor-derived mBDOs and mPDOs. On stage 1, organoids were cultured in the mEM (without noggin) supplemented with BMP7 which was used to initiate endocrine fate; On stage 2, organoids were cultured in the pancreatic differentiation media (mDM9) containing vascular endothelial growth factor (VEGF), exendin 4, Triiodothyronine (T3), A8301, DAPT, Heparin, activin A and retinoic acid (RA), to enhance pancreatic differentiation (Figure 3A). The results showed that the organoids treated with BMP7 for 7 d were significantly up-regulated the expression of neurogenin 3 (*Ngn3*) which is an endocrine progenitor marker (Figure 3B); after the second stage of differentiation culture, the organoids were dramatically increased the mRNA expression of pancreatic β-cell specific gene Insulin2 (*Ins2*), also that of the other pancreatic endocrine and exocrine genes, including Glucagon (*GCG*), Somatostatin (*SST*), *and* Amylase (*AMY*) demonstrated by qRT-PCR analysis (Figure 3C). Immunofluorescence staining confirmed that both endocrine and exocrine cells, including C-peptide (C-PEP), GCG, SST and AMY positive cells could be detected in the organoids (Figure 3D and S3). Flow cytometry analysis showed that the ratio of insulin-producing cells (C-PEP^+^) in the differentiated mBDOs and mPDOs were 8.49 ± 0.45% and 7.23 ± 0.63%, respectively (Figure 3E and 3F). All the above-mentioned results indicate that Trop2^+^ mBDOs have a similar pancreatic differentiation potential as mPDOs do, including endocrine cells or exocrine cells.

**Figure 3.**
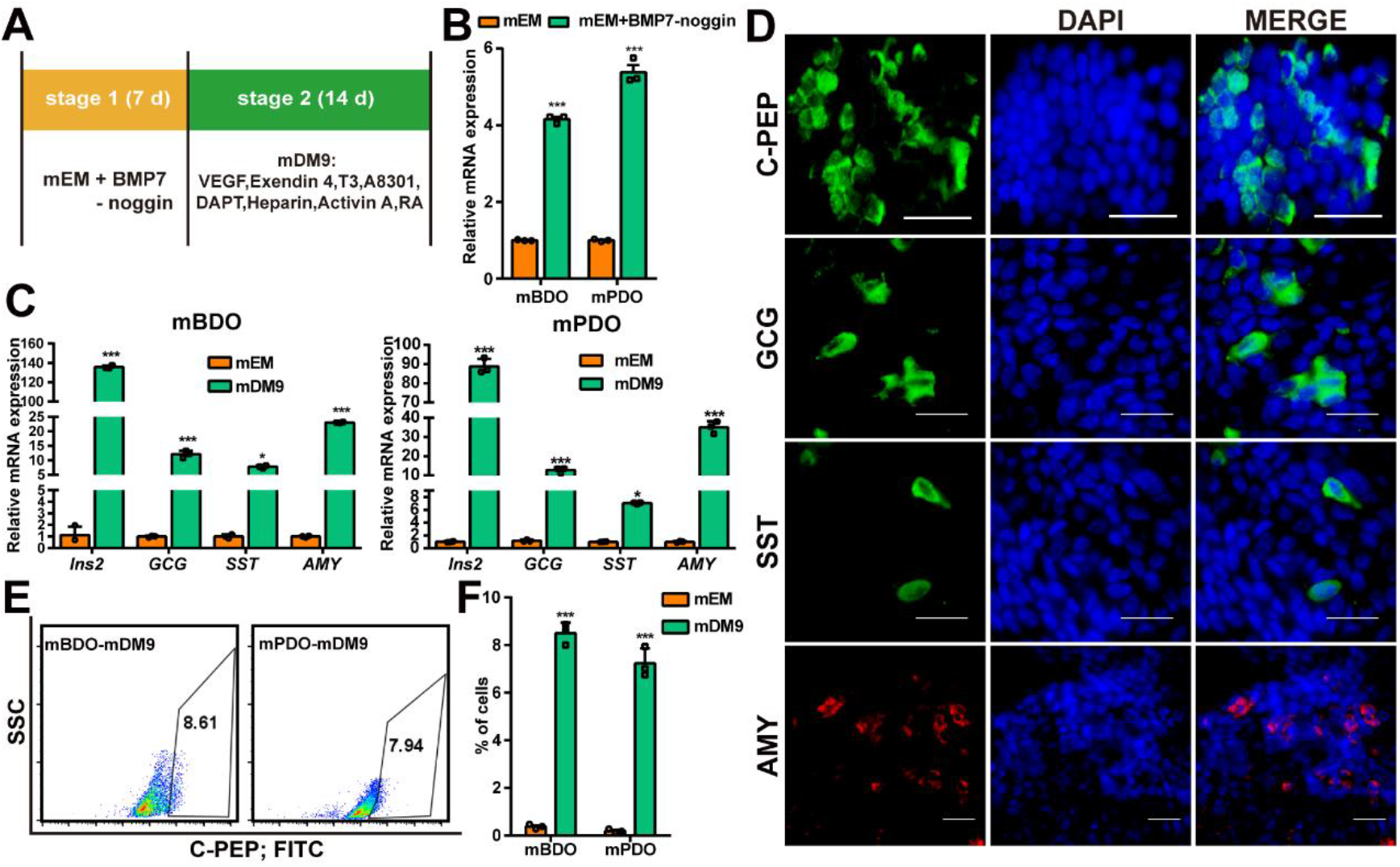
mBDOs and mPDOs have similar differentiation capability. **(A)** Schematic of organoids induced differentiation into pancreatic lineage using BMP7 and mDM9. **(B)** qRT-PCR showing the relative fold changes of *Ngn3* mRNA levels of mBDOs and mPDOs cultured with mEM or mEM (without noggin) containing BMP7. **(C)** The relative expression changes of *Ins2, GCG, SST* and *AMY* mRNA levels under indicated culture condition detected by qRT-PCR. **(D)** Immunostaining of C-PEP, GCG, SST and AMY in the differentiated mBDOs. Scale bar, 50 μm. **(E)** Representative image of FCM plot of the differentiated mBDOs and mPDOs stained with C-PEP. **(F)** The percentages of C-PEP^+^ cells in the differentiated mBDOs and mPDOs were detected by FCM. n = 3 independent cultures. Data are presented as mean ± SD. *p <0.05, ***p <0.001

### TLY142 increases the ratio of insulin-secreting cells differentiated from mBDOs

Compared with the pancreatic ductal cells, biliary ductal cells are able to obtain from patients themselves or the donors through minimally invasive laparoscopic surgery, thus they have a broader application prospect in the future (Sampaziotis et al., 2017). Therefore, we used mBDOs for the further experiments. To increase the efficiency of mBDOs differentiation into pancreatic β-cells, we attempted to find the potential chemical small molecules in chemdiv 160M compounds (https://www.chemdiv.com/). We first screened the possible activators of glucagon like peptide 1 (GLP 1) by computer-aided analysis, and then verified them using a GLP-1R reporter cell line. 18 chemical compounds were filtered out (Table S1). We found that TLY142, one of the 18 chemical molecules, could significantly promote mBDOs expressing insulin 2 after supplemented it into the mDM9 media (Figure 4A and 4B). To further test the differentiation effect of TLY142, we performed qRT-PCR and found that the expression of endocrine genes (*Ins2, GCG*, and *SST*) were increased (Figure 4C). Immunofluorescence staining demonstrated an increased proportion of C-peptide^+^ cells after the addition of TLY142, but not that of GCG, and SST in organoids (Figure 4D). Flow cytometry analysis showed that, compared with the mBDOs differentiated in mDM9 (Figure 3E), TLY142 supplement significantly increased the ratio of C-peptide^+^ cells from 8.49 ± 0.45% up to 19.90 ± 0.62% (Figure 4E). Glucose-stimulated C-peptide secretion tests showed that the differentiated mBDOs with mDM9 plus TLY142 had a glucose-response ability, and could increase the intracellular calcium content after the high glucose and potassium chloride stimulation (Figure 4G). The above results demonstrate that the chemical molecule TLY142 could remarkably promote the production of insulin-secreting cells from mBDOs, which present a glucose-responsive capacity.

**Figure 4.**
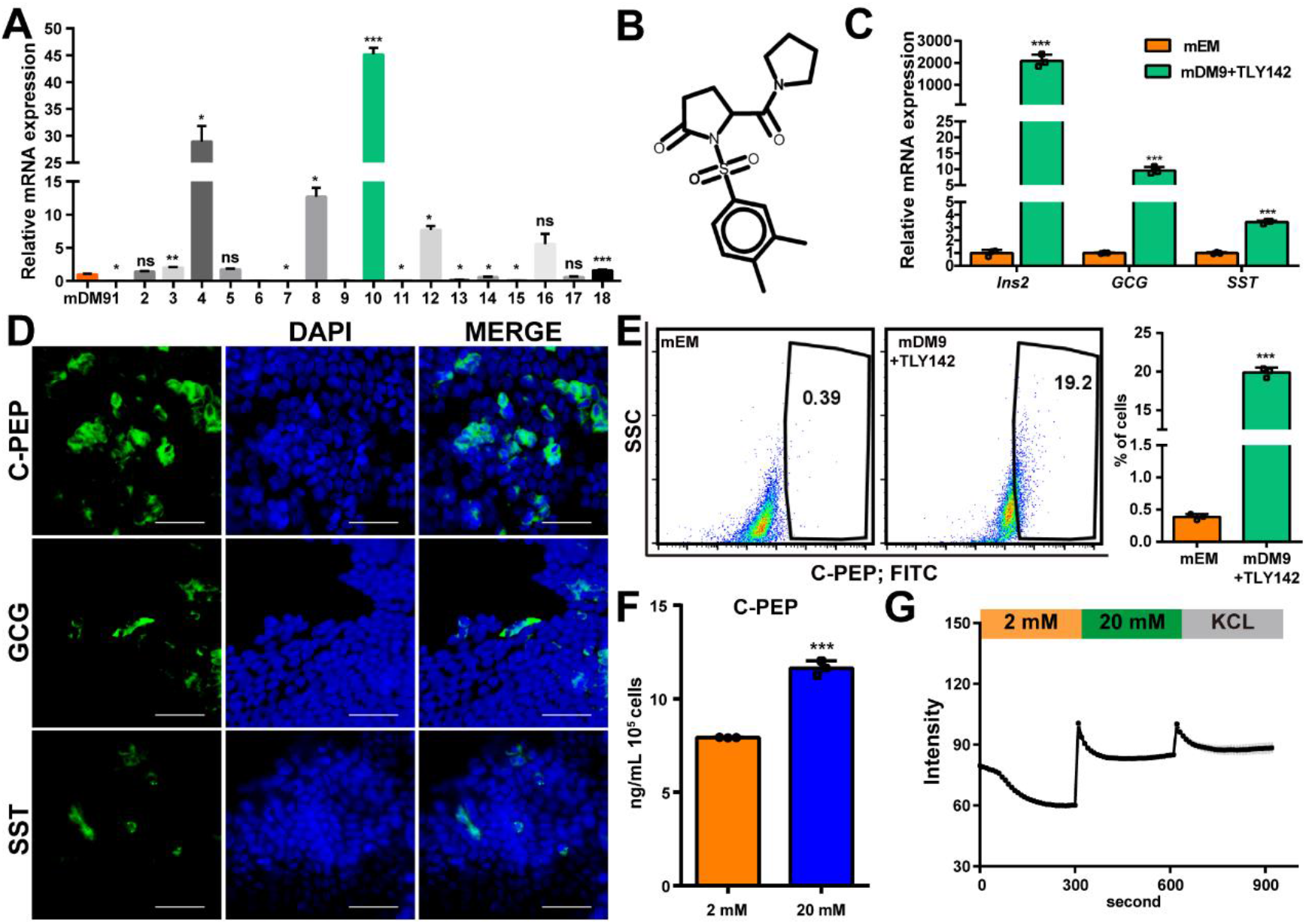
TLY142 enhances the differentiation of organoids into glucose-responsive insulin-secreting cells. **(A)** The relative fold changes of *Ins2* mRNA levels in mBDOs cultured with different small molecule compounds based on mDM9 medium. **(B)** Structural formula of TLY142 compound. **(C)** qRT-PCR showing the relative fold changes of *Ins2, GCG*, and *SST* mRNA levels in mBDOs under indicated culture condition. **(D)** Representative immunofluorescence images show the expression of C-PEP, GCG and SST in the differentiated mBDOs under TLY142 plus condition. Scale bar, 50 μm. **(E)** Representative image of FACS plot of mBDOs under the indicated culture condition stained with C-PEP. The percentage of C-PEP^+^ cells in the differentiated mBDOs were counted by FCM. **(F)** ELISA analysis of C-peptide secretion from the differentiated mBDOs under TLY142 condition challenged with 2 mM and 20 mM glucose, with a 10-min incubation for each concentration. **(G)** Representative examples of calcium signaling traces from the differentiated mBDOs under TLY142 condition when challenged sequentially with 2 mM, 20 mM glucose, and 30 mM KCl. The X axis represents time. n = 3 independent cultures. Data are presented as mean ± SD. *p <0.05, **p <0.01, ***p <0.001, ns, not significant.

### Transplantation of differentiated mBDOs by mDM9 plus TLY142 exerts a significantly hypoglycemic effect in STZ-induced diabetic mice

To further examine the function of differentiated mBDOs, we transplanted the mBDOs treated with mDM9 or mDM9 plus TLY142 under the kidney capsules of diabetic nude mice which were induced by streptozotocin (STZ). One week after transplantation, comparing to the STZ group, the blood glucose began to decrease in TLY142 plus treated organoid-transplantation group mice, whereas the mDM9-only treated organoids-transplanted mice maintains a high level. After 2 weeks of transplantation, the mice receiving the TLY142 plus treated mBDOs had a more stable hypoglycemic effect than mDM9 (Figure 5A). The intraperitoneal glucose tolerance tests (IPGTT) showed a significant improvement in diabetic mice receiving TLY142 plus treated organoids compared with those receiving only mDM9 treated control (Figure 5B and 5C). Interestingly, compared with the mDM9 treated organoids-transplanted mice, TLY142 plus treated organoid-transplanted mice presented more significant glucose tolerance, which was similar as islet transplanted mice. Meanwhile, serum insulin level was significantly increased after transplantation of TLY142 plus treated mBDOs (Figure 5D). After removing the transplanted kidneys, a sharp rise in blood glucose was found in the TLY142 plus treated organoid-transplanted mice (Figure 5A), which proving that the previous recovery of blood glucose in diabetic mice was indeed caused by the transplanted organoids. Immunostaining of the removed kidney showed the presence of Insulin^+^ and GCG^+^ cells (Figure 5E), indicating that the transplanted mBDOs could indeed differentiate into pancreatic endocrine cells. The static analysis of the stained sections revealed that 36.09 ± 4.92% of the transplanted cells were Insulin^+^. Overall, TLY142 plus mDM9 treated mBDOs had a significant hypoglycemic effect in STZ-induced diabetic mice.

**Figure 5.**
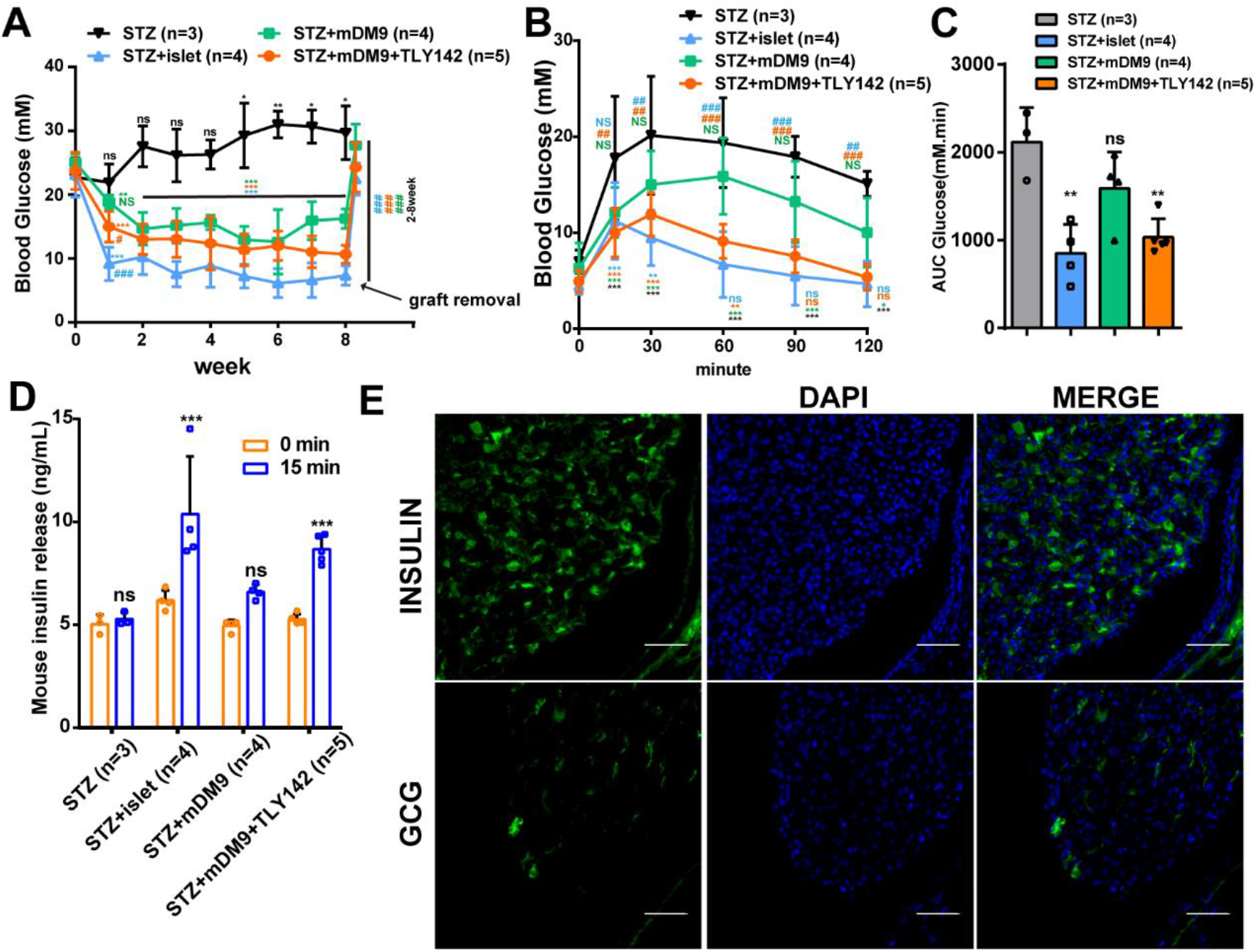
The differentiated mBDOs with TLY142 have a better hypoglycemic effect *in vivo*. **(A)** Transplantation of the differentiated mBDOs lowers the blood glucose in STZ-induced diabetic mice. Grafted mBDOs were removed via nephrectomy at 8 weeks after transpalantation. **(B)** and **(C)**. Blood glucose and area under curves (AUCs) changes after IPGTT in mice transplanted with each indicated cells. Data are presented as mean ± SD. Two-way ANOVA with Sidak’s test is used for comparison of multiple groups. Within a group, each time point is compared with t = 0 level, *p <0.05, **p <0.01, ***p <0.001, ns, not significant; Between groups, each group is compared with the STZ group, ^#^p <0.05, ^##^p <0.01, ^###^p <0.001. NS, not significant. Numbers of mice measured per group are indicated in brackets. **(D)** ELISA analysis of insulin level in each indicated group before and after (15 min) glucose injection. **(E)** The production of insulin and glucagon in the engrafted mBDOs underneath the kidney capsule after two months were detected by immunofluorescence staining.

### Trop2^+^ progenitor-derived hBDOs have multiple differentiation potential

To investigate whether Trop2 can mark the progenitor cells in human biliary ducts, we performed immunostaining on the human extrahepatic bile duct (hEBD) sections, and found that Trop2 was also highly expressed on the epithelium of hEBD (Figure S4A). Flow cytometry analysis further demonstrated that a majority of epithelial cells in hEBD were Trop2^+^ (Figure 6A), which was consistent with the previous results of mEBD. We performed organoid formation assay on Trop2^+^ cells isolated from hEBD by flow cytometry (Figure 6A), and found that 12.80 ± 3.12% of them could form typical lumen-like organoids (Figure 6B and S4B), which expressed the progenitor marker (SOX9, PDX1), the pancreatic development related genes (HNF1B, HNF4A), and the epithelium-specific genes (EpCAM, E-cadherin) (Figure S4C). The above results indicate that Trop2 can also be used as a marker for enrichment of the progenitor cells with organoid formation capacity from hEBD.

**Figure 6.**
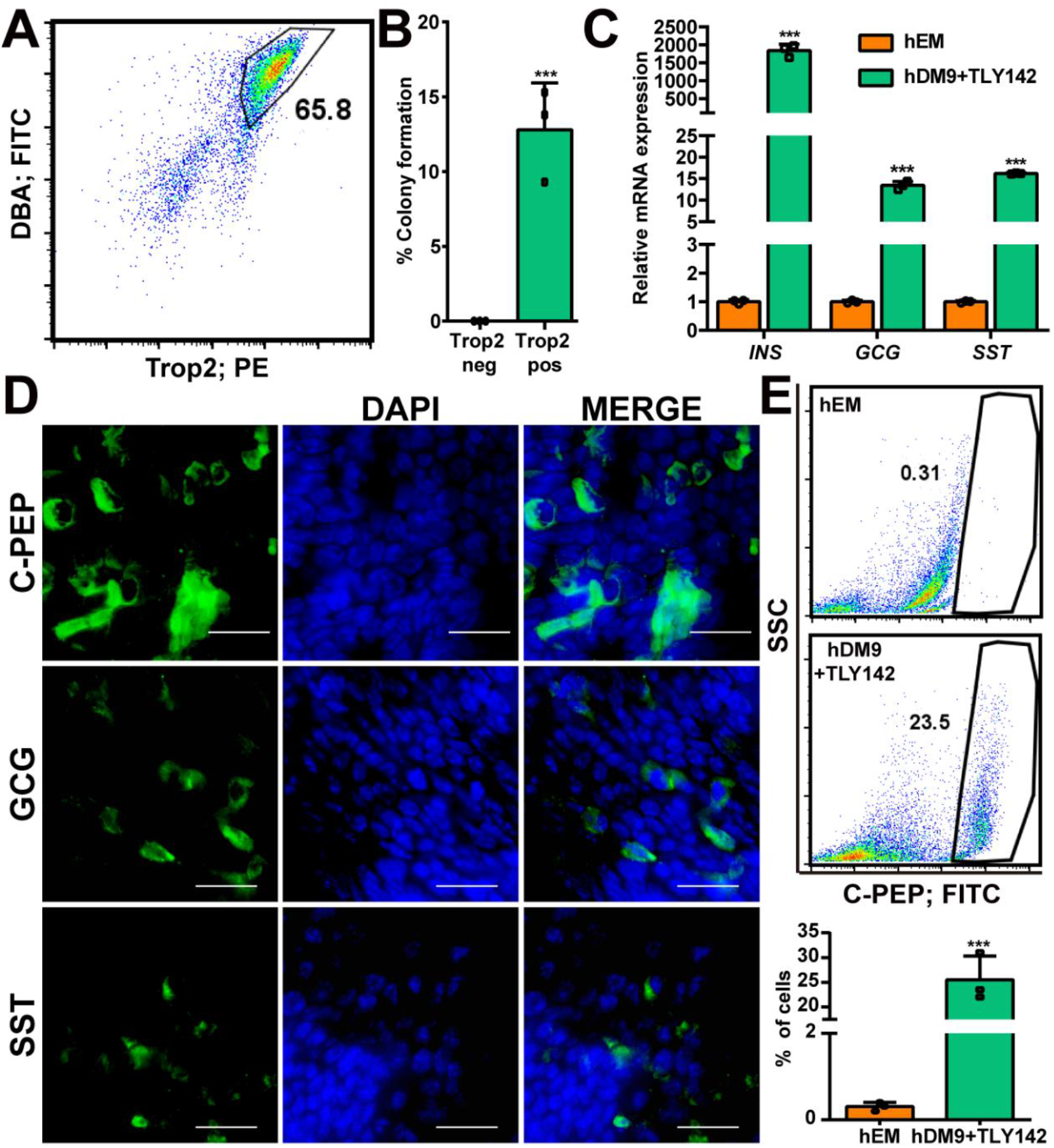
Trop2^+^ progenitors in hEBD formed hBDOs can be differentiated into endocrine cells promoted by TLY142 *in vitro*. **(A)** Using Trop2 and DBA as markers to isolate progenitors from hEBD via FCM. **(B)** The organoid formation efficiency of the Trop2^+^ DBA^+^ fractions. neg, negative, pos, positive. n=3, 1000 cells were seeded each well. **(C)** qRT-PCR assays show the relative fold changes of *INS, GCG*, and *SST* mRNA levels in hBDOs differentiated with TLY142. **(D)** Representative immunofluorescence images show the expression of C-PEP, GCG and SST in the differentiated hBDOs under TLY142 condition. Scale bar, 50 μm. **(E)** Representative image of FCM plot of hBDOs under each indicated culture condition stained with C-PEP. The percentage of C-PEP^+^ cells were counted in the differentiated hBDOs detected by FCM. n=3, independent cultures. ***p <0.001

Next, we tried to determine whether hBDOs could be differentiated into pancreatic endocrine cells. We further optimize the human pancreatic differentiation medium (hDM9) containing TLY142. qRT-PCR results showed that the differentiated hBDOs significantly increased the expression of endocrine genes, including β cells, α cells, and δ cells compared with the undifferentiated ones (Figure 6C). Immunofluorescence staining demonstrated a high ratio of C-peptide-positive cells, also a small proportion of GCG-and SST-positive cells existed (Figure 6D). Flow cytometry defined the ratio of C-peptide-positive in the differentiated hBDOs was 25.50 ± 4.82% (Figure 6E). These results indicate that Trop2^+^ hBDOs are multipotential and could be used to produce a relative proportion of insulin-secreting cells *in vitro*.

### Trop2^+^ hBDO-derived β cells were functional both *in vitro* and *in vivo*

To verify whether the insulin-producing cells differentiated from hBDOs were functional, we conducted *in vitro* and *in vivo* physiological experiments on the differentiated hBDOs by hDM9 plus TLY142. Glucose-stimulated C-peptide secretion tests showed that the differentiated hBDOs responded to a high concentration of glucose (20 mM) (Figure 7A), and potassium chloride to present a physiological calcium wave showed by Fluo4-AM staining (Figure 7B). In addition, we transplanted the differentiated hBDOs treatment with hDM9 plus TLY142 into STZ-induced diabetic nude mice. After 2 weeks post transplantation, blood glucose levels in the transplanted mice were significantly reduced and maintained continuously (Figure 7C). The *in vivo* glucose stimulation revealed a robust secretion of human insulin in transplanted diabetic mice (Figure 7D). At the same time, mice that received differentiated hBDOs exhibited a strong glucose clearance ability (Figure 7E and 7F). Removal of the transplanted kidney by nephrectomy resulted in a dramatic increase of blood glucose (Figure 7C), which confirmed that the functional insulin-secreting cells resided in the kidney transplants. Analysis of the removed kidneys revealed that insulin- and glucagon-producing endocrine cell types were detectable, with the vast majority being INS^+^ β-cells (45.68 ± 13.31%) (Figure 7G). Above all, Trop2^+^ hBDO-derived insulin-secreting cells were functional both *in vitro* and *in vivo*.

**Figure 7.**
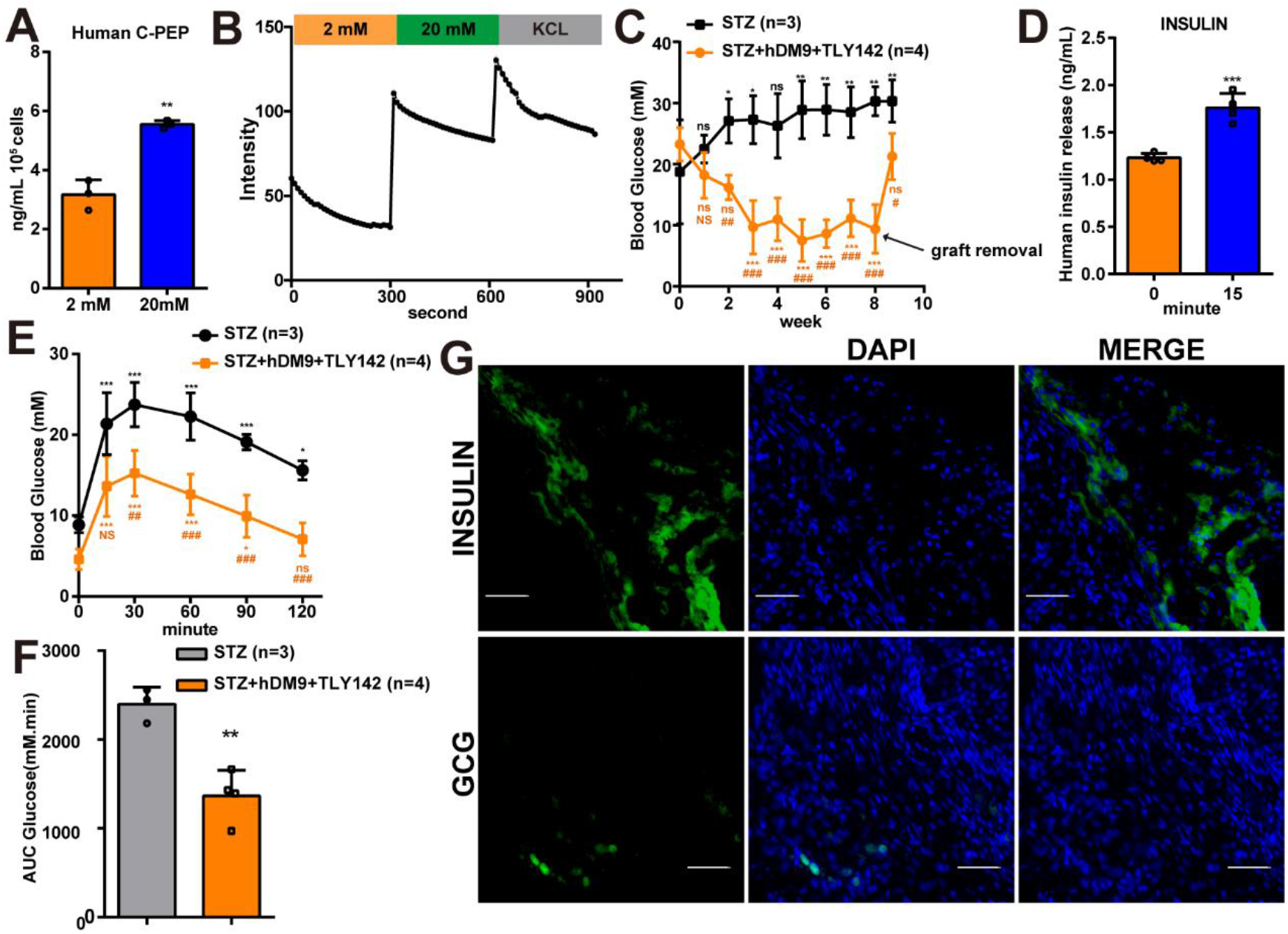
Differentiated hBDOs with TLY142 have glucose response ability both *in vitro* and *in vivo*. **(A)** ELISA analysis of C-peptide levels of hBDOs differentiated under TLY142 condition challenged with 2 mM and 20 mM glucose, with a 10-min incubation for each concentration. **(B)** Calcium signaling traces from differentiated hBDOs under TLY142 condition when challenged sequentially with 2 mM, 20 mM glucose, and 30 mM KCl. The X axis represents time. **(C)** Blood glucose levels in STZ-treated mice after transplantation with hBDOs differentiated under TLY142 condition. Graft removal: nephrectomy to remove transplanted cells. Numbers of mice measured per group are indicated in brackets. Data are presented as mean ± SD. Two-way ANOVA with Sidak’s test is used for comparison of multiple groups. Within a group, each time point is compared with t = 0 level, *p <0.05, **p <0.01, ***p <0.001, ns, not significant; Between groups, each group is compared with the STZ group, ^#^p <0.05, ^##^p <0.01, ^###^p <0.001. NS, not significant. **(D)** ELISA analysis of human insulin levels in mice after transplantation with the differentiated hBDOs before and after (15 min) glucose injection. Numbers of mice measured per group are indicated in brackets. **(E)** and **(F)** Blood glucose and area under curves (AUCs) changes after IPGTT in mice transplanted with the differentiated hBDOs. **(G)** Immunofluorescence image shows the expression of insulin and glucagon in the differentiated hBDOs engrafted underneath the kidney capsule after two months. Data are presented as mean ± SD.

## Discussion

In this study, we used Trop2 as a novel marker to identify a population of progenitors with the long-term organoid forming capacity from the mouse extrahepatic bile ducts as well as the pancreatic ducts, and demonstrated that they had similar self-renewal and multi-potential differentiation capacity. We further showed that BMP7 was an effective cytokine to convert ductal progenitor to a multipotent state, thus it was crucial for endocrine differentiation. By screening chemical molecules, we found that TLY142 greatly increased the efficiencies of mBDOs differentiation into insulin-secreting cell based on BMP7 initiation. Moreover, we proved that Trop2^+^ human extrahepatic bile duct progenitors are long-term organoid-forming and multipotential, TLY142 enhanced hBDOs differentiation into insulin-secreting cells at a high proportion, and transplantation of the differentiated mBDOs and hBDOs *in vivo* could efficiently response to glucose challenge and cure the high glucose of diabetic mice.

Our findings extend the usage of biliary duct epithelia as the potential beta-cell source in cell replacement therapy of diabetes. As revealed in recent findings that the progenitors isolated from autologous biliary ducts have been demonstrated to be organoid-forming, and capable of repairing the damaged biliary ducts through feedback transplantation (Sampaziotis and Muraro, 2021). We and others demonstrated that biliary ductal progenitors could be isolated using markers, such as Trop2 and DBA, from biliary ductal epithelia, whereas these epithelia are easier to be acquired from autologous biliary duct via microsurgery, or donated biliary ducts (Sampaziotis et al., 2017). Our studies further demonstrated that the biliary ductal organoids presented similar pancreatic endocrine potential as those pancreatic ductal organoids, especially insulin-secreting cell differentiation (Figure 4D and S3). Transplantation of the differentiated human BDOs effectively cured diabetes of model diabetic mice, indicating biliary ductal progenitor-derived organoids are a liable source in cell transplantation to treat diabetes.

Trop2 is a useful marker for enrichment of hepatic-biliary-pancreatic progenitor cells. In liver, Trop2 is not expressed in the normal liver but expressed specifically in oval cells after liver injury (Okabe et al., 2009). A human liver cell atlas reveals that Trop2^+^ progenitor population with strong potential to form bipotent liver organoids (Aizarani et al., 2019), establishing it serves as a marker to distinguish hepatic progenitor cells from normal parenchymal liver cells. In this study, we first verify that Trop2 represents a subpopulation of progenitor cells in biliary and pancreatic ducts capable of forming organoids and multi-potential. Of note, a previous report shows that Trop2 is expressed in the luminal epithelium of mouse and human extrahepatic bile duct exclusively (Matsui et al., 2018). However, we found that Trop2 was also expressed in the gland epithelia surrounding pancreatic ducts and biliary ducts. The subtle difference on the Trop2 positive localization in biliary ducts might due to the distinct sensitivity in different studies.

A high ratio of glucose-responsive beta cells differentiated from ductal organoids provide a feasible beta-cell source in cell transplantation therapy of diabetes. Both BMP7 and chemical small molecule TLY142 are crucial in augmentation of beta-cell differentiation from organoids. We found that BMP7 was important for the transition of ductal organoid to Ngn3^+^ endocrine progenitor state, which was consistent with the previous report that BMP7 significantly increased Ngn3 expression (Klein et al., 2015). As a result, after BMP7 withdraw, we defined a DM9 media, which significantly increased the ratio of insulin-secreting cells to 7-8% of total cells, rather higher than no BMP7 addition control (1-5%) (Huch et al., 2013; Loomans et al., 2018). It has been reported that Ngn3^+^ progenitor cells are multipotential, and could be differentiated into multiple pancreatic lineages (Gradwohl et al., 2000), however, the non-islet differentiation weakened the production of beta cells. Here, we dig out a chemical molecule named TLY142, which significantly promoted the differentiation of beta cells from 7% to over 20% *in vitro* and about 40% *in vivo*. Furthermore, the differentiated insulin-secreting cells could response to glucose challenge both *in vitro* and *in vivo*. The increased percentage of insulin-secreting cells from mouse or human BDOs assure the hypoglycemic effect after transplantation under the kidney capsule.

### Limitations of study

We have demonstrated that transplantation of the differentiated mBDOs or hBDOs could efficiently lower the high blood glucose in diabetic mice. However, the proportion of insulin-secreting cells in differentiated BDOs still required to be increased, since it is still lower than that of beta cells in islets. Further mechanical exploration and small molecule mining which aims to enhance the efficiency of beta-cell differentiation will promote the usage of autonomous or heterogenous biliary duct organoids in the cell replace therapy of diabetes.

## Acknowlegement

This work was supported by the National Natural Science Foundation of China (No. 32072801 and 81972724). The Fundamental Research Funds for the Central Universities of China (2572020AW36).

## Author Contributions

Miao Liu, Wen Yu, and Chun-Bo Teng designed, performed and analyzed most experiments; Yannan Wang, Bingru Yan, Jie Shang, and Huan Wu performed *in vivo* experiment and analysis. Sheng Tai and Congyi Zhang performed resection and collection of human extrahepatic bile duct tissues, and help with tissue section detection. Liang Jin provided preliminary screening of small molecule libraries and prepared the manuscript together with Miao Liu, Wen Yu, and Chun-Bo Teng.

## Declaration of interests

The authors declare no competing interests.

## Data and code availability

All high-throughput sequencing data, both raw and processed files, have been deposited in NCBI’s Gene Expression Omnibus and are accessible under the accession number GEO: GSE196394. To view the sequencing data, please go to https://www.ncbi.nlm.nih.gov/geo/query/acc.cgi?acc=GSE196394 and enter token wvifgcsaptmhpsn into the box.

## MATERIALS AND METHODS

### Animals and human biliary tissues

NU/NU nude immunodeficient mice (3 months of age, 25-30 g) and C57BL/6N mice (2 months of age, 20-25 g) were purchased from Beijing Charles River Laboratory Animal Technology Co., Ltd., and maintained in the SPF-grade laboratory animal room in College of Life Sciences, Northeast Forestry University. The mouse feeding conditions strictly followed the requirements of the SPF-grade laboratory animal room. The protocol for use of animals was approved by the Animal Care and Ethics Committee of Northeast Forestry University, and all procedures were carried out in accordance with the approved guidelines.

We collected human extrahepatic bile ducts from the donating patients in the Second Affiliated Hospital of Harbin Medical University. All patients who participated in this study provided informed consent. This study was approved by the Research Ethics Committee.

### Preparation of organoids from mouse extrahepatic bile duct and pancreatic duct

Mouse extrahepatic bile duct (mEBD) and pancreatic duct (mPD) were obtained from C57BL/6N mouse by direct dissection under a dissecting microscope (Agbunag et al., 2006), and then were digested at 37°C with collagenase IV for 20 min after mechanical mincing, washed and centrifuged to resuspend the cells in Matrigel (Corning), and 600 μL mouse expansion media (mEM) was added to 24 wells. All cells were cultured in a humidified incubator with 95% air and 5% CO_2_ at 37°C. Freshly prepared media can be stored at 4 degrees for 2 weeks. The main components of mEM included D/F12 basal media, 1% Glutamax (Hyclone), 1% penicillin-streptomycin (Hyclone), 2% B27 Supplement (minus VA, Thermo), 30% Wnt3A condition media, 1 mM N-acetylcysteine (Sigma), 50 ng/mL EGF (MCE), 100 ng/mL R-Spondin-1 (RSPO1, R&D), 25 ng/mL Noggin (Peprotech), 100 ng/mL FGF10 (Peprotech), 10 mM Nicotinamide (Sigma) and 10 μM Y27632 (Sigma). When the luminal organoid obtained, the culture media was changed into mEM without Wnt3A and Y27632. Media were changed every other day and organoids were passaged at a ratio of 1:3-1:5 every 7 to 8 days.

### Preparation of human extrahepatic bile duct organoids

Take the donated human extrahepatic bile duct samples for preparation of human extrahepatic bile duct organoid (hBDOs). Briefly, the samples were placed in a 10-cm dish containing sterile PBS, longitudinally cut open the catheter with dissecting scissors, and scraped the catheter surface with a sterile glass slide. The flushing fluid was collected after flushing the catheter with sterile PBS and were digested into single cells with Trypsin. The Trop2^+^DBA^+^ cells were collected by FCM analysis, sorted and seeded for organoid growing. Briefly, the sorted cells were resuspended in Matrigel, and left at 37°C for 10 min. After solidification, human expansion media (hEM) consist of mEM supplemented with 1% N2 supplement (Thermo), 10 nm Gastrin (Sigma), 3 μM Prostaglandin E2 (PGE2, MCE) and 5 μM A8301 (MCE) was added and cultured at 37°C with 5% CO_2_. When the cells proliferated into cystic structures, they were switched to hEM media without Wnt3A and Y27632. The media needed to be replaced every other day. The organoids were passaged 2 times every week at a ratio of 1:3-1:5.

### RNA-sequencing and analysis

Organoids from a 24 well plate were harvested in 1 mL of TRIzol reagent. All used organoid lines were at passage 3 or 4 after established. Freshly obtained mEBD and mPD were minced mechanically, and resuspend in TRIzol reagent. The total RNA of each sample was sent to ANOROAD GENOME (Beijing, China) for mRNA sequencing on the Illumina HiSeq X Ten platform. All data are available in the NCBI’s Gene Expression Omnibus. DEGSeq2 (Anders and Huber, 2010) was used for differential gene expression analysis. Differential gene screening mainly refers to the difference fold (Fold change value) and adjust p value (<0.05) as relevant indicators.

### Chemical compounds

The chemical structure of TLY142 was given in Figure 4B. TLY142 IUPAC name is 1-(3, 4-dimethylphenyl) sulfonyl–5-(pyrrolidine–1-carbonyl) pyrrolidine–2-one (https://pubchem.ncbi.nlm.nih.gov/compound/53008166), which used in the animal and in vitro studies was purchased from chemhui.

### Differentiation of PDOs/BDOs into Pancreatic cell lineages

The mPDOs or mBDOs with good growth status were cultured with mEM (without noggin) and 100 ng/mL BMP7 for 7 days after passage, and then replaced with fresh mDM9 with or without 1μM TLY142 for another 14 days. The media were changed every other day. After the end of differentiation, RNA samples and immunostained samples were collected for identification, and the differentiation efficiencies were detected by flow cytometry. The main components of mDM9 include high-glucose DMEM media, 1% Glutamax (Hyclone), 1% penicillin-streptomycin (Hyclone), 2% B27 Supplement (Thermo), 1% ITS (Thermo), 50 ng/mL VEGF (Peprotech), 50 ng/mL Exendin 4 (MCE), 25 ng/mL Noggin (Peprotech), 750 ng/mL RSPO1 (R&D), 1 μM T3 (MCE), 10 μM A8301 (MCE), 5 μM DAPT (MCE), 10 μg/mL heparin (MCE), 100 ng/mL Activin A (Peprotech), 20 ng/mL FGF2 (MCE), 4.4 mM Nicotinamide (Sigma) and 1 μM Retinoic acid (MCE). For hBDOs, the pancreatic differentiation was induced in hDM9 as a similar step as mice. Human pancreatic β cell differentiation medium (hDM9) is based on mDM9 with 1% N2 Supplement.

### Extraction of total RNA and synthesis of cDNA

Total RNA was extracted using the TRIzol method. Reverse transcription of RNA into cDNA was performed according to the instructions of the Vazyme ^™^ HiScript ® II Q Select RT SuperMix Kit (Vazyme). The synthesized cDNA was aliquoted and stored at -80°C for quantitative Real-time PCR (qRT-PCR).

### qRT-PCR

The primers were presented in Table 1. For the reaction system and conditions, refer to the instructions for use of ChamQ Universal SYBR qPCR Master Mix Kit (Vazyme), qRT-PCR was performed by Roche Light Cycle 480 fluorescent quantitative PCR instrument. The analysis was performed using the Light Cycler 480 built-in software analysis module, Ct values were calculated by Abs Quant/2nd Derivative Max, 3 replicates were set for each samples and β-actin was used as the reference gene to calculate the relative expression level of target gene by 2^-△△Ct^ method.

**Table 1.**
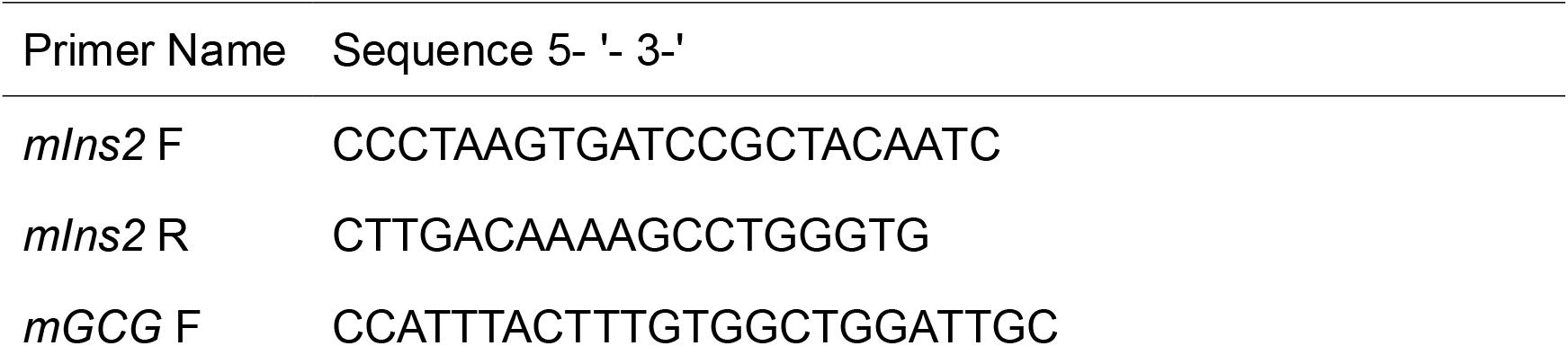
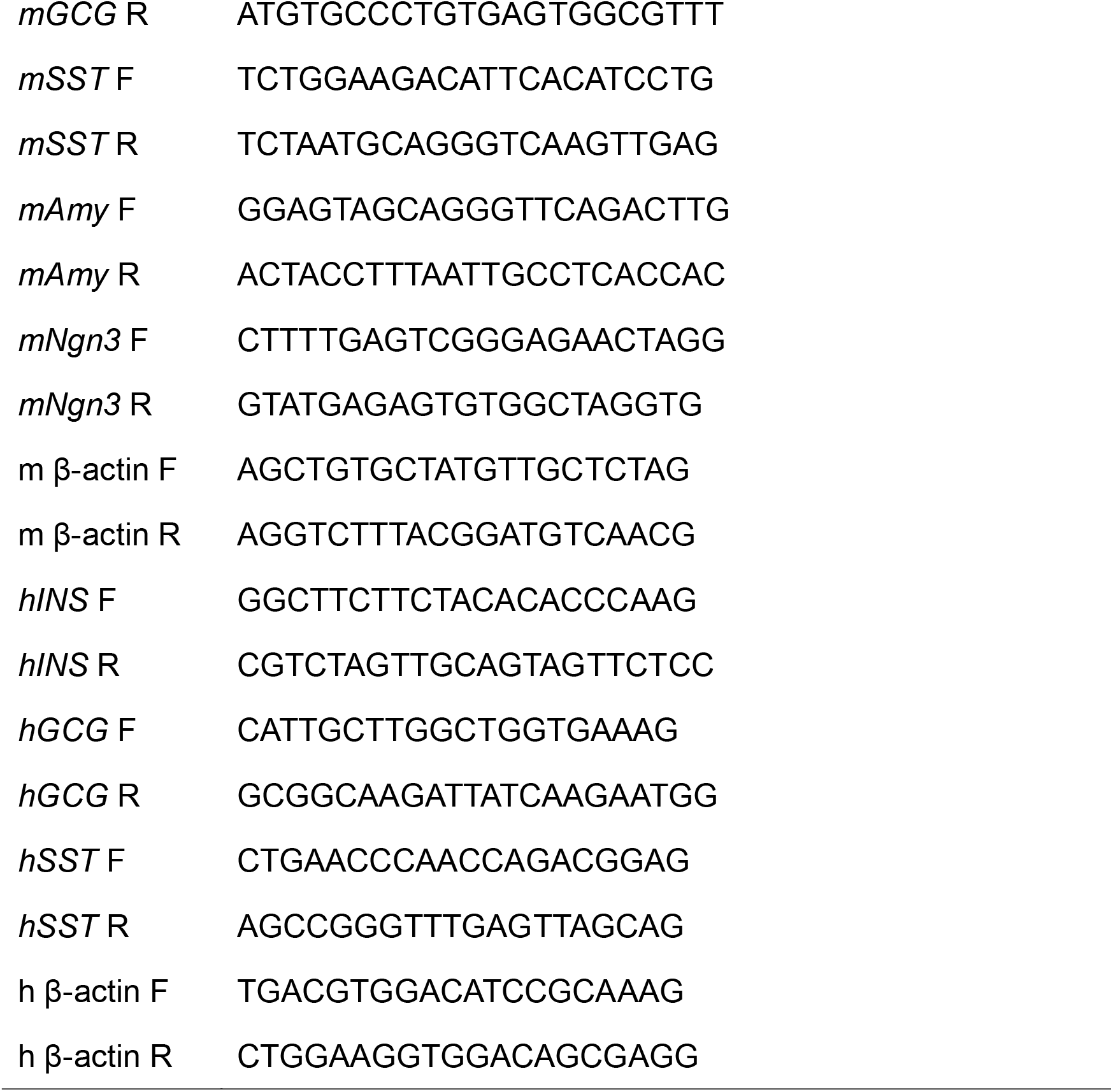
Primer sequences for qRT-PCR.

### Cryosections and patches of organoids

Cryosection preparation: Tissue or cultured organoids were fixed in 4% PFA for 15 min at room temperature. The tissue or organoids were then dehydrated in 30% sucrose overnight at 4°C and then embedded in OCT to prepare approximately 10 μm sections using a thermostated cryostat.

Preparation of organoid patch: The cultured organoids were digested by dispase to get rid of Matrigel, and then were fixed in 4% PFA and washed with cold PBS to ensure the clearance of Matrigel. During the cleaning process, the organoids could naturally settle overnight at 4°C to ensure the recovery rate of organoids. The recovered organoids were resuspended with PBS solution, dropped on glass slides, placed at 37°C for drying, and dried for immunofluorescence staining.

### Immunofluorescence staining

Prepared sections or patches were punched with 0.3% TritonX 100. 10% horse serum (Cunninghamia lanceolata) was incubated for 1 h at 37 for blocking. Primary antibodies (Table 2) were incubated overnight at 4°C. Secondary antibody incubated for 2 h at room temperature in the dark. Hoechst 33342 (Beyotime) was used for nuclear staining. After mounting using anti-fluorescence burst mounting media, the stained samples were observed by Delta vision, a high-resolution cell imager. Immunostaining of frozen sections from the resected kidney requires the use of Vector® TrueVIEW® Autofluorescence Quenching Kit (Vectorlabs) to remove the autofluorescence.

**Table 2.**
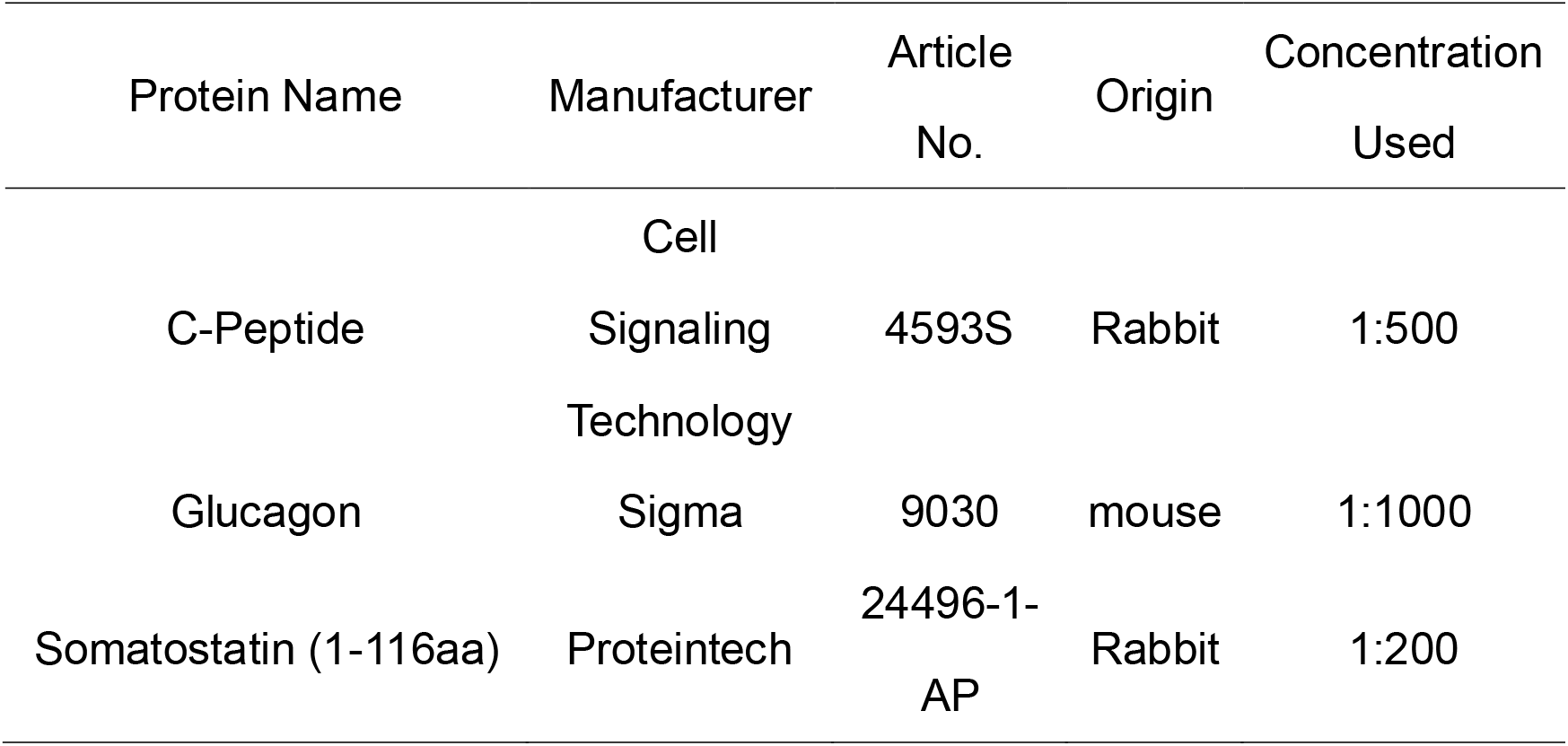
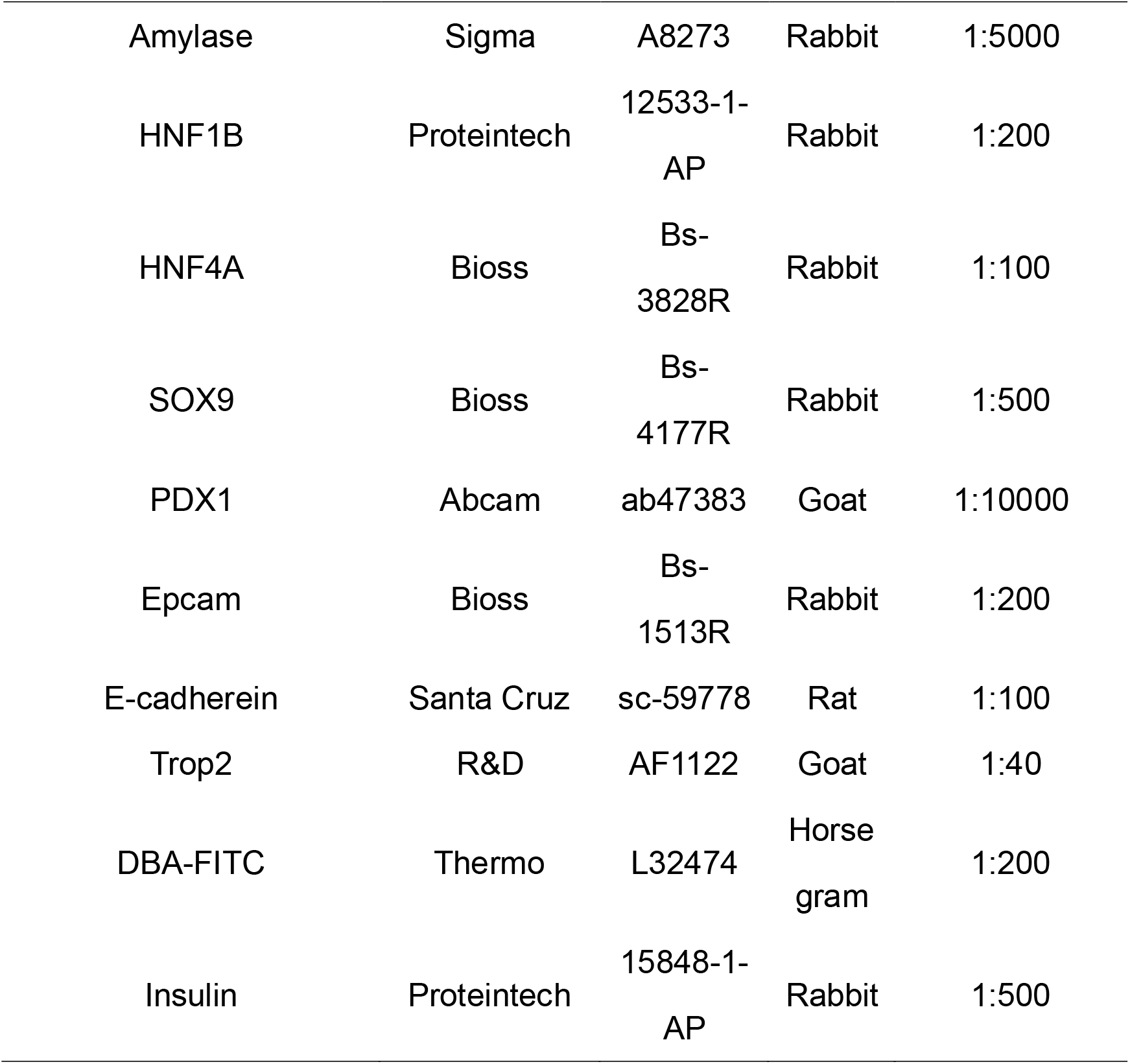
Primary antibodies used in immunofluorescence.

### Flow cytometry – sorting and analyzing

The cultured organoids were blown using precooled high-glucose DMEM medium and centrifuged to remove Matrigel, which could be repeated to ensure the clearance of Matrigel. Organoids or scratched epithelia were digested into single cells using 0.25% of trypsin. At least 5×10^5^ cells were taken from each group of experiments. Cells were fixed on ice using 4% PFA, punched with 0.1% TritonX-100, and blocked with 5% BSA. Primary antibody diluted in 1% BSA was incubated on ice for 40 min, and isotype antibody diluent was used as control. The Secondary antibody (1:100) diluted in 1% BSA and incubated for 30 min on ice in the dark. After washing the cells with PBS, the cells were resuspended with PBS and loaded for detection or sorting.

### Glucose-stimulated C-peptide secretion

Differentiated organoids depleted of Matrigel were resuspended in sugar-free Krebs solution, washed two to three times, and placed in low-adsorption culture plates overnight. The following day, Krebs solution containing 2 mM glucose was added for resuspension of organoids after removal by centrifugation of sugar-free Krebs. After incubation for 10 min, the supernatant was collected by centrifugation. Cells were washed with sugar-free Krebs followed by resuspension with 20 mM glucose in Krebs and incubated for 10 min, the supernatant was collected by centrifugation. C-peptide levels in supernatant samples were analyzed using a mouse/human C-peptide ELISA kit (Mlbio, ml-1015542) following standard protocols.

### Calcium ion detection after differentiation

The differentiated organoids were harvested after dispase dissociation and cultured adherently for 24 h in a 35 mm laser confocal dish without Matrigel. Fluo4-AM was used to incubate the samples for 30 min. Then, Fluo4-AM was washed away with Krebs buffer, and the cells were incubated in Krebs buffer at 37°C for an additional 15 min. The cells were then detected under a high-resolution cell imager, Delta vision, for the acquisition of high-resolution time-series images. Time series images were acquired every 10 s. The progression of glucose challenge and stimulation time during imaging were as follows: low glucose (2 mM) for 5 min, Krebs wash, high glucose (20 mM) for 5 min, Krebs wash, and a final incubation of 2 mM glucose containing 30 mM potassium chloride (KCl) for 5 min. For the measurement of mean fluorescence intensity, images were quantified by image J software.

### Animal model

Nude mice were injected with STZ at a concentration of 160 mg/kg, and blood glucose levels were measured 72 hours later. The mice with Blood glucose more than 16.8 mmol/L for 3 consecutive days were used for renal subcapsular transplantation surgery, and then were monitored blood glucose.

### Transplantation of organoids under the capsules of diabetic mice kidney

After differentiation culture, the Matrigel around the organoids was removed using precooled high-glucose DMEM media. The organoids were collected after centrifugation for subcapsular renal transplantation. Approximately 10^6^ cells or 500 islets were transplanted per mouse, and organoids were resuspended using 30% Matrigel for transplantation. Eight weeks after transplantation, the kidneys with transplanted cells were enucleated, fixed by PFA, dehydrated in sucrose, and subjected to frozen section and immunostaining. The islets of mouse were obtained from C57BL/6N mice using the previously reported method (Zmuda et al., 2011).

### IPGTT glucose tolerance test

For intraperitoneal glucose tolerance test (IPGTT), mice were fasted for 16 h with free access to water. Baseline blood samples were collected from the tail of fully conscious mice, followed by intraperitoneal injection of D-glucose (2 g/kg body weight), and blood glucose was monitored using a Roche glucometer by tail vein blood samples at 15, 30, 60, 90, and 120 min after glucose administration. The area under the curve (AUC) of the IPGTT was determined using the trapezoidal rule. Serum samples before and after glucose injection were also collected and measured by ELISA.

### Statistical analysis

Each experiment was performed at least three sets of repetitions. Analyses were performed using a two-way analysis of variance (ANOVA) to compare more than two treatments and were followed by a Sidak’s test for multi-group comparisons with a single control group, as applicable. * P <0.05, ** P <0.01, ***P <0.001 were considered statistically significant.

## Supplementary figure legend

**Figure S1**

**(A)** Serial images of the growing and passaged mPDOs and mBDOs. Scale bar, 50 μm.

**(B)** and **(C)**. mPDOs and mBDOs were stained with the primary antibodies against E-cadherin, HNF1B, HNF4A. Scale bar, 50 μm.

**Figure S2**

**(A)** Immunofluorescence images show the expression of Trop2 in mPD and mEBD epithelium (DBA^+^). Scale bar, 50 μm.

**(B)** Immunofluorescence staining of Trop2 in pancreatic acinar cell (Amylase^+^) and islet (Insulin^+^). Scale bar, 50 μm.

**(C)** Representative image of mPDOs or mBDOs grown out of the sorted Trop2^+^DBA^+^ cells from mPD or mEBD by flow cytometry for 14 days after seeding at a 10^3^ density each well.

**(D)** Immunofluorescence staining of Trop2 in mPDOs and mBDOs produced by the sorted Trop2^+^DBA^+^ cells. Scale bar, 50 μm.

**(E)** FCM plot of Trop2 and DBA staining for mPDOs and mBDOs grown from the sorted Trop2^+^DBA^+^ cells.

**Figure S3**

Representative immunofluorescence images show the expression of C-PEP, GCG and SST in the differentiated mPDOs under mDM9 plus TLY142 condition. Scale bar, 50 μm.

**Figure S4**

**(A)** Immunofluorescence images show the expression of Trop2 in hEBD epithelium (DBA^+^). Scale bar, 100 μm.

**(B)** Representative image of hBDOs grown out of the sorted Trop2^+^DBA^+^ cells from hEBD by flow cytometry at the 5^th^, the 20^th^ days and the tenth passage after seeding at a 10^3^ density each well.

**(C)** hBDOs were stained with the primary antibodies against EPCAM, SOX9, PDX1, E-cadherin, HNF1B, HNF4A. Scale bar, 50 μm.

## Notes

### Competing Interest Statement

The authors have declared no competing interest.

