## Supplementary figures and images for "A chemical molecule promotes Trop2^+^ biliary duct organoids differentiation into insulin-secreting cells"

### supplement figure 1

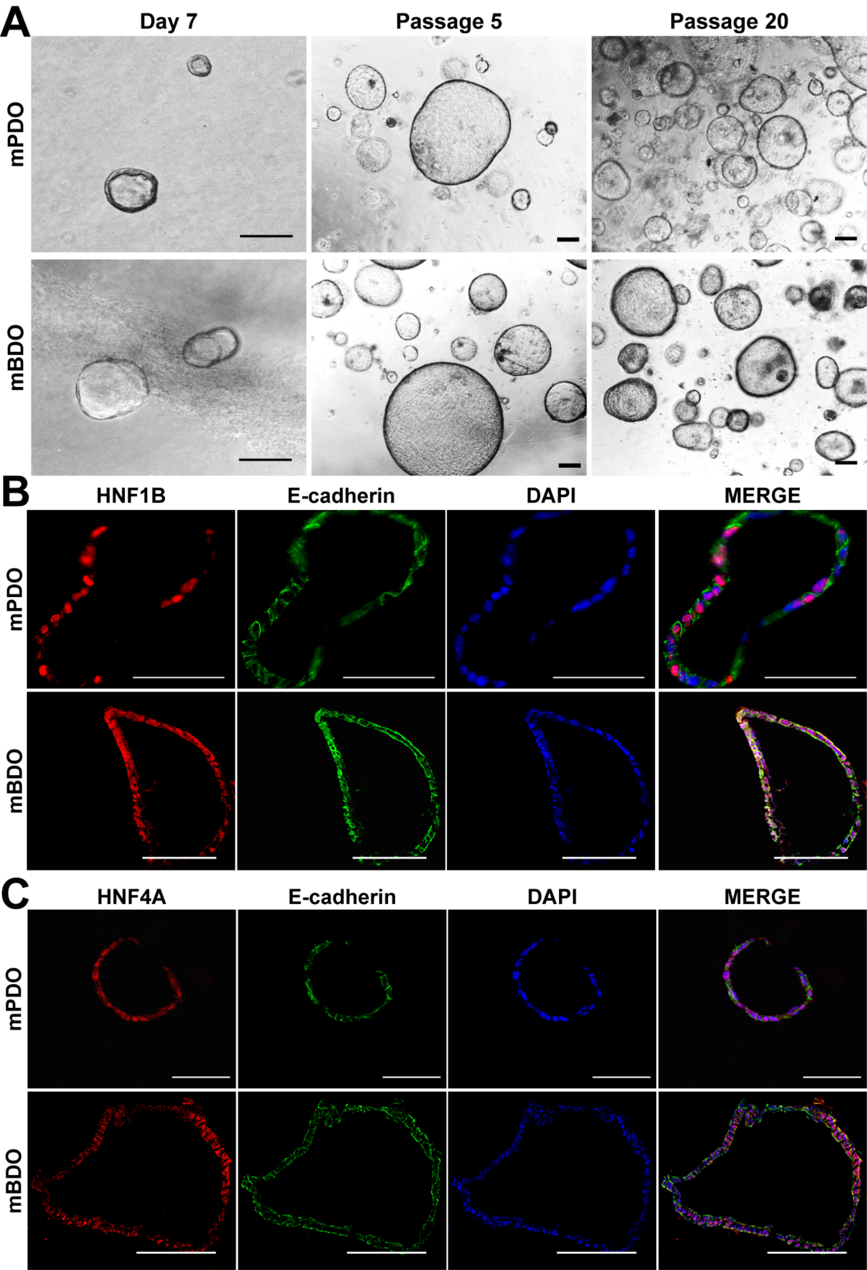

### supplement figure 2

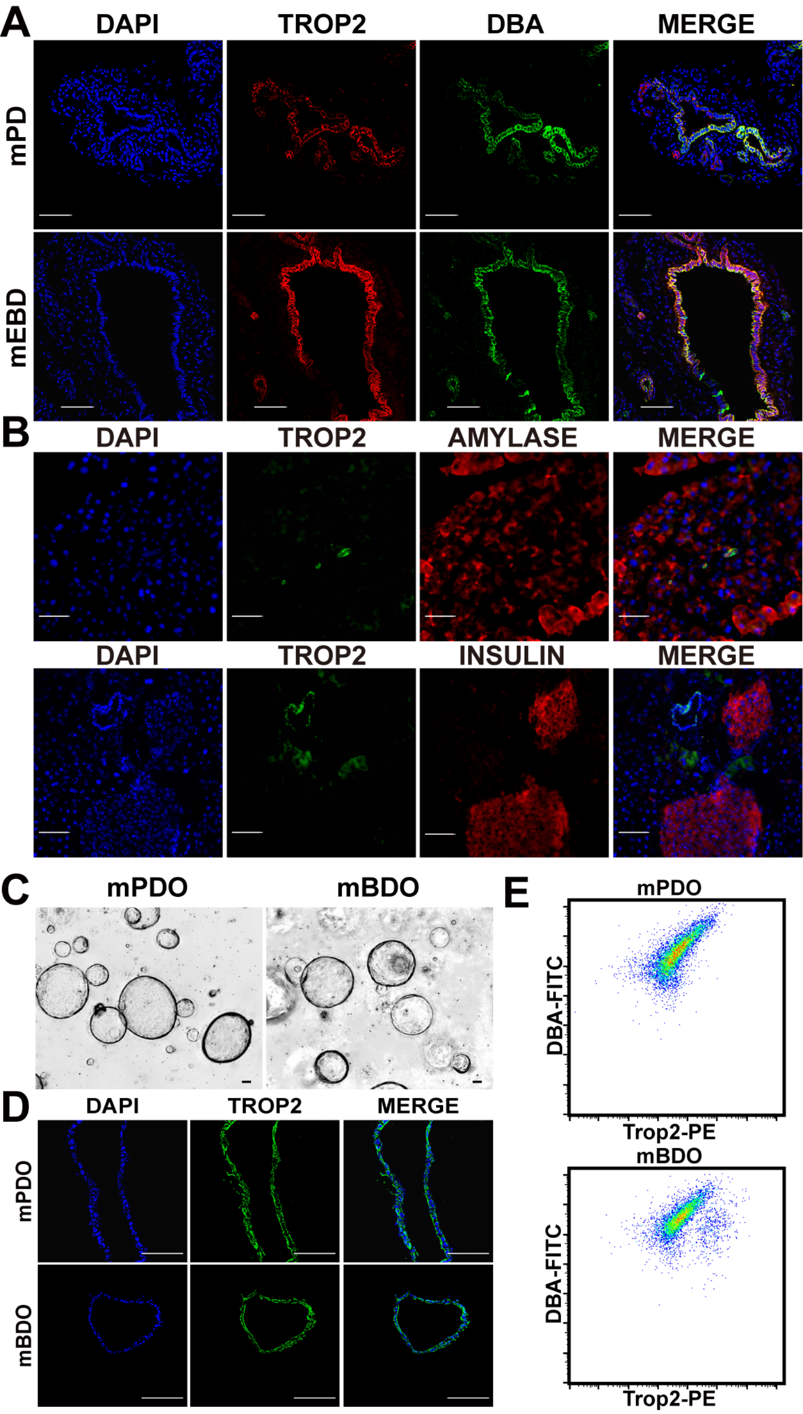

### supplement figure 3

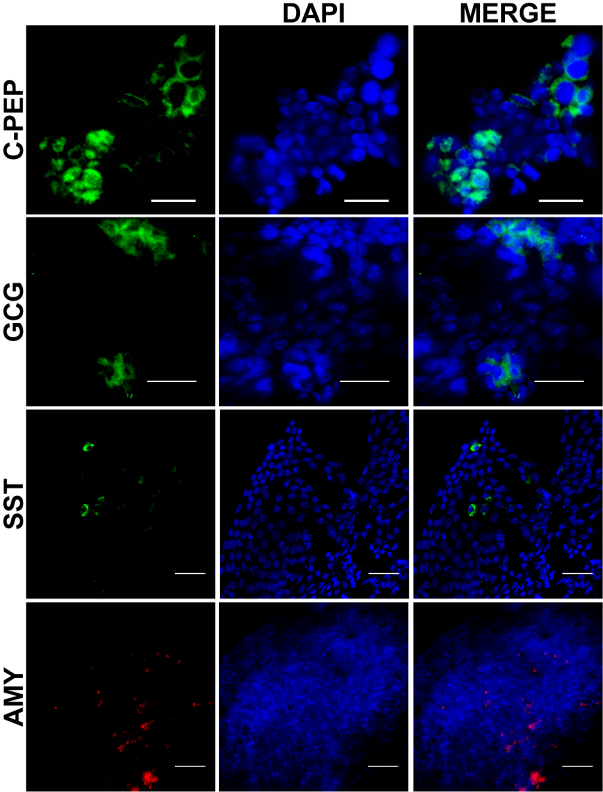

### supplement figure 4

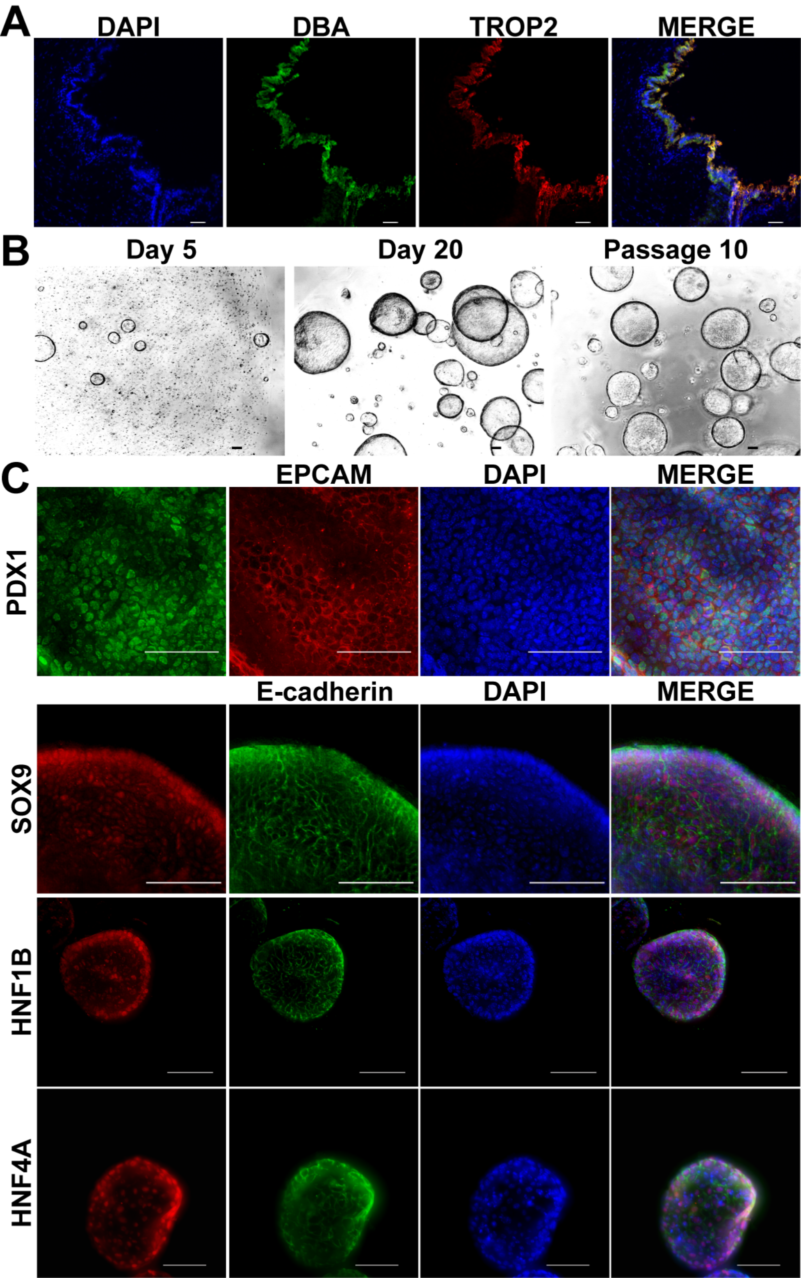
