## supplement table 1 for "A chemical molecule promotes Trop2^+^ biliary duct organoids differentiation into insulin-secreting cells"

| | Structure | IUPAC Name | |
|---|---|---|---|
| 1 | 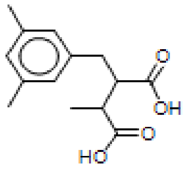 | 2-(3,5-Dimethyl-benzyl)-3-methyl-succinic acid | https://pubchem.ncbi.nlm.nih.gov/substance/328101137 |
| 2 | 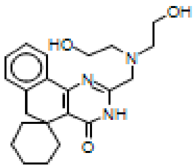 | 2-{[bis(2-hydroxyethyl)amino]methyl}-3H-spiro[benzo[h]quinazoline-5,1'-cyclohexan]-4(6H)-one | https://pubchem.ncbi.nlm.nih.gov/compound/135449533 |
| 3 | 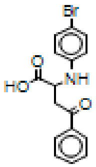 | 2-[(4-bromophenyl)amino]-4-oxo-4-phenylbutanoic acid | https://pubchem.ncbi.nlm.nih.gov/substance/328238220 |
| 4 | 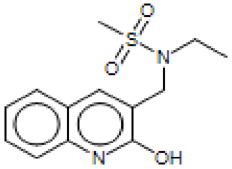 | N-ethyl-N-[(2-hydroxyquinolin-3-yl)methyl]methanesulfonamide | https://pubchem.ncbi.nlm.nih.gov/substance/328489909 |
| 5 | 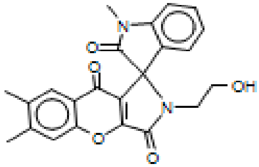 | 2-(2-hydroxyethyl)-1',6,7-trimethyl-2H-spiro[chromeno[2,3-c]pyrrole-1,3'-indole]-2',3,9(1'H)-trione | https://pubchem.ncbi.nlm.nih.gov/compound/16013149 |
| 6 | 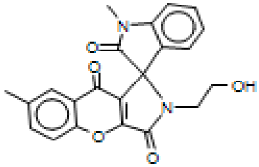 | 2-(2-hydroxyethyl)-1',7-dimethyl-2H-spiro[chromeno[2,3-c]pyrrole-1,3'-indole]-2',3,9(1'H)-trione | https://pubchem.ncbi.nlm.nih.gov/substance/328492655 |

| | | | |
|---|---|---|---|
| 7 | 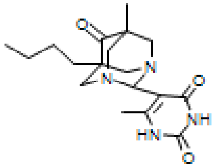 | 5-(5-butyl-7-methyl-6-oxo-1,3-diazatricyclo[3.3.1.1~3,7~]dec-2-yl)-6-methylpyrimidine-2,4(1H,3H)-dione | https://pubchem.ncbi.nlm.nih.gov/compound/16014026 |
| 8 | 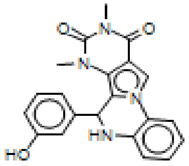 | 6-(3-hydroxyphenyl)-7,9-dimethyl-6,7-dihydropyrimido[4',5':3,4]pyrrolo[1,2-a]quinoxaline-8,10(5H,9H)-dione | https://pubchem.ncbi.nlm.nih.gov/substance/328540065 |
| 9 | 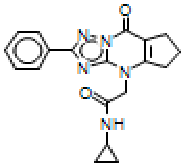 | N-cyclopropyl-2-(8-oxo-2-phenyl-5,6,7,8-tetrahydro-4H-cyclopenta[d][1,2,4]triazolo[1,5-a]pyrimidin-4-yl)acetamide | https://pubchem.ncbi.nlm.nih.gov/substance/328578701 |
| 10 | 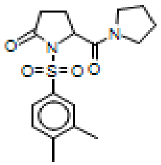 | 1-(3,4-Dimethylphenyl)sulfonyl-5-(pyrrolidine-1-carbonyl)pyrrolidin-2-one | https://pubchem.ncbi.nlm.nih.gov/compound/53008166 |
| 11 | 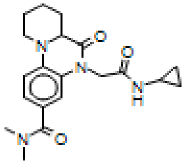 | 5-[2-(cyclopropylamino)-2-oxoethyl]-N,N-dimethyl-6-oxo-6,6a,7,8,9,10-hexahydro-5H-pyrido[1,2-a]quinoxaline-3-carboxamide | https://pubchem.ncbi.nlm.nih.gov/substance/328876029 |
| 12 | 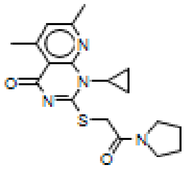 | 1-cyclopropyl-5,7-dimethyl-2-[(2-oxo-2-pyrrolidin-1-ylethyl)thio]pyrido[2,3-d]pyrimidin-4(1H)-one | https://pubchem.ncbi.nlm.nih.gov/substance/328886576 |

| | | | |
|---|---|---|---|
| 13 | 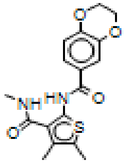 | N-{4,5-dimethyl-3-[(methylamino)carbonyl]-2-thienyl}-2,3-dihydro-1,4-benzodioxine-6-carboxamide | https://pubchem.ncbi.nlm.nih.gov/substance/328894048 |
| 14 | 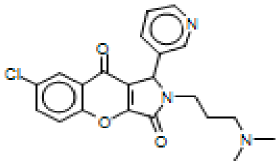 | 7-chloro-2-[3-(dimethylamino)propyl]-1-pyridin-3-yl-1,2-dihydrochromeno[2,3-c]pyrrole-3,9-dione | https://pubchem.ncbi.nlm.nih.gov/substance/328914132 |
| 15 | 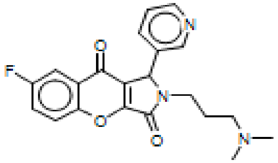 | 2-[3-(dimethylamino)propyl]-7-fluoro-1-pyridin-3-yl-1,2-dihydrochromeno[2,3-c]pyrrole-3,9-dione | https://pubchem.ncbi.nlm.nih.gov/substance/328914151 |
| 16 | 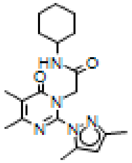 | N-cyclohexyl-2-[2-(3,5-dimethyl-1H-pyrazol-1-yl)-4,5-dimethyl-6-oxopyrimidin-1(6H)-yl]acetamide | https://pubchem.ncbi.nlm.nih.gov/substance/328917944 |
| 17 | 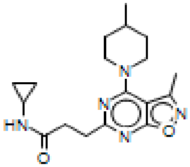 | CHEMDIV NAME: L553-1335.sdf-chemdiv_compunds | |
| 18 | 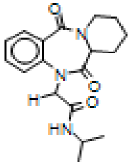 | CHEMDIV NAME: L560-0018.sdf-chemdiv_compunds | |
